# All-optical interrogation of neural circuits in behaving mice

**DOI:** 10.1101/2021.06.29.450430

**Authors:** Lloyd E. Russell, Henry W. P. Dalgleish, Rebecca Nutbrown, Oliver M. Gauld, Dustin Herrmann, Mehmet Fişek, Adam M. Packer, Michael Häusser

## Abstract

Recent advances combining two-photon calcium imaging and two-photon optogenetics with digital holography now allow us to read and write neural activity *in vivo* at cellular resolution with millisecond temporal precision. Such “all-optical” techniques enable experimenters to probe the impact of functionally defined neurons on neural circuit function and behavioural output with new levels of precision. This protocol describes the experimental strategy and workflow for successful completion of typical all-optical interrogation experiments in awake, behaving head-fixed mice. We describe modular procedures for the setup and calibration of an all-optical system, the preparation of an indicator and opsin-expressing and task-performing animal, the characterization of functional and photostimulation responses and the design and implementation of an all-optical experiment. We discuss optimizations for efficiently selecting and targeting neuronal ensembles for photostimulation sequences, as well as generating photostimulation response maps from the imaging data that can be used to examine the impact of photostimulation on the local circuit. We demonstrate the utility of this strategy using all-optical experiments in three different brain areas – barrel cortex, visual cortex and hippocampus – using different experimental setups. This approach can in principle be adapted to any brain area for all-optical interrogation experiments to probe functional connectivity in neural circuits and for investigating the relationship between neural circuit activity and behaviour.

## INTRODUCTION

A fundamental goal in neuroscience is to understand how the brain encodes information in patterns of neural activity that can be used to guide behaviour. Addressing this challenge requires methods that allow for the controlled manipulation of neuronal activity in vivo to determine which features of neural activity are most relevant to the behaviour (Jacobs *et al*., 2009; Panzeri *et al*., 2017) - such as the spike rate (London *et al*., 2010; Histed and Maunsell, 2014), spike timing (Panzeri *et al*., 2001; Gollisch and Meister, 2008; Shusterman *et al*., 2011; Doron *et al*., 2014), spike number (Huber *et al*., 2008; Doron *et al*., 2014), as well as the functional identity (Harris and Mrsic-Flogel, 2013; Pinto and Dan, 2015) and spatial distribution of the neurons active during behaviour. These questions can now be addressed using a recently introduced “all-optical” experimental strategy, which combines two-photon calcium imaging (Denk *et al*., 1990, 1994; Tian *et al*., 2009; Grienberger and Konnerth, 2012; Chen *et al*., 2013) with two-photon optogenetics (Boyden *et al*., 2005; J. P. Rickgauer and Tank, 2009; Packer *et al*., 2012; Prakash *et al*., 2012) and digital holography (Nikolenko *et al*., 2008; Papagiakoumou *et al*., 2008, 2010). This approach allows simultaneous reading and writing of neural activity *in vivo* (**Figure 1**), and has been made possible by the combined effort of many labs who have developed elements of the all-optical toolkit, and combined them to achieve successful implementations of the strategy (Szabo *et al*., 2014; Rickgauer *et al*., 2014; Packer *et al*., 2015; Carrillo-Reid *et al*., 2016; Shemesh *et al*., 2017; Pégard *et al*., 2017; Zhang *et al*., 2018; Forli *et al*., 2018; Mardinly *et al*., 2018; Chettih and Harvey, 2019; Jennings *et al*., 2019; Marshel *et al*., 2019; Gill *et al*., 2020). This strategy has already been used for a wide range of experiments, including mapping functional connectivity of circuits (Chettih and Harvey, 2019; Jennings *et al*., 2019; Marshel *et al*., 2019; Russell *et al*., 2019; Dalgleish *et al*., 2020; Daie *et al*., 2021), and modulation of behaviour through the targeted manipulation of functionally defined neurons in several brain areas (Carrillo-Reid *et al*., 2019; Jennings *et al*., 2019; Marshel *et al*., 2019; Russell *et al*., 2019; Dalgleish *et al*., 2020; Gill *et al*., 2020; Robinson *et al*., 2020; Daie *et al*., 2021). As all-optical interrogation becomes more widely used, it is crucial that the potential pitfalls and limitations of the approach are recognised, and that rigorous standards are set by the field to facilitate the implementation and interpretation of experiments using this approach.

**Figure 1.**
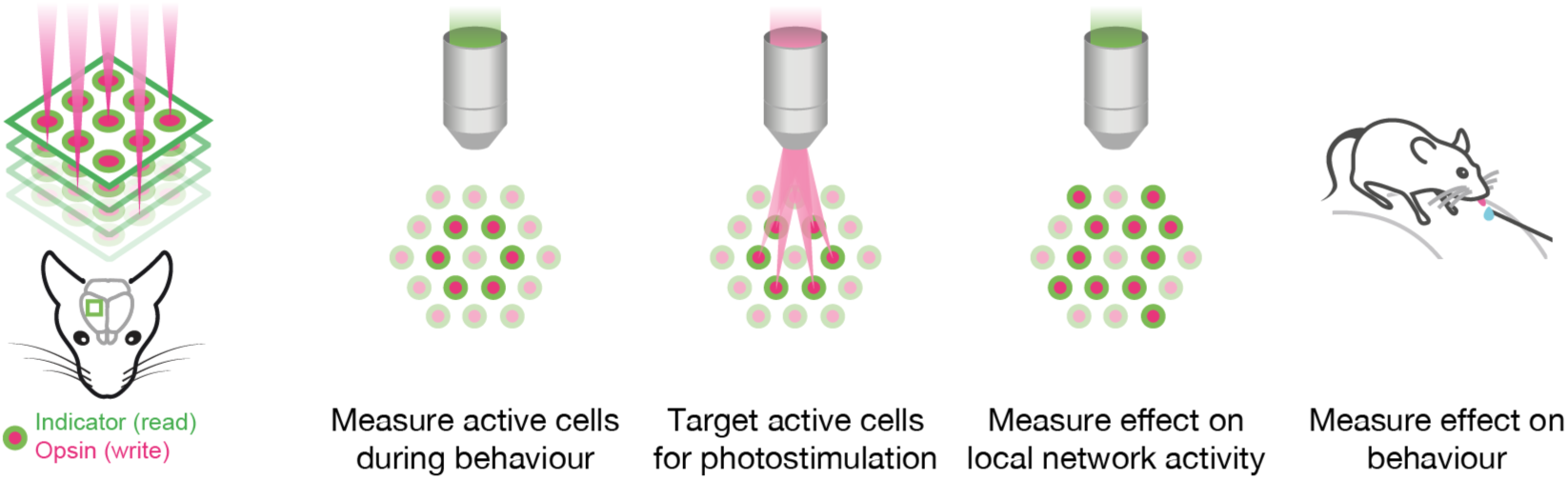
Conceptual goals of all-optical interrogation experiments. Schematic diagram illustrating the basic elements of all-optical interrogation studies, showing the typical sequence used in an experiment.

The all-optical approach is challenging, regardless of the specific implementation, as it requires many complex experimental steps, as well as the co-ordinated interaction between multiple software and hardware modules (**Figure 2, 3**). For example, all-optical systems involve two lightpaths, one for imaging and one for photostimulation. These lightpaths each comprise laser sources, power modulation devices, beam steering mirrors and on the photostimulation side beam patterning devices to enable multiple neurons to be stimulated at once. The imaging lightpath may employ volumetric scanning with a variety of methods (electrically tunable lenses, piezo elements, spatial light modulators) to enable larger populations to be recorded and thus targeted by the photostimulation pathway (Mardinly *et al*., 2018; Yang *et al*., 2018; Marshel *et al*., 2019; Russell *et al*., 2019). These two lightpaths must be calibrated such that they are co-registered, allowing the photostimulation laser to hit precise locations in the imaging FOV. To perform experiments the photostimulation is targeted to neurons that have typically been identified by some anatomical or functional property. These targeted neurons, in order to be stimulated, must coexpress two proteins enabling their activity to be read out as well as manipulated. Finally, the stimulation of these neurons could be triggered by an external event, perhaps as part of a behavioural task the animal is performing.

**Figure 2.**
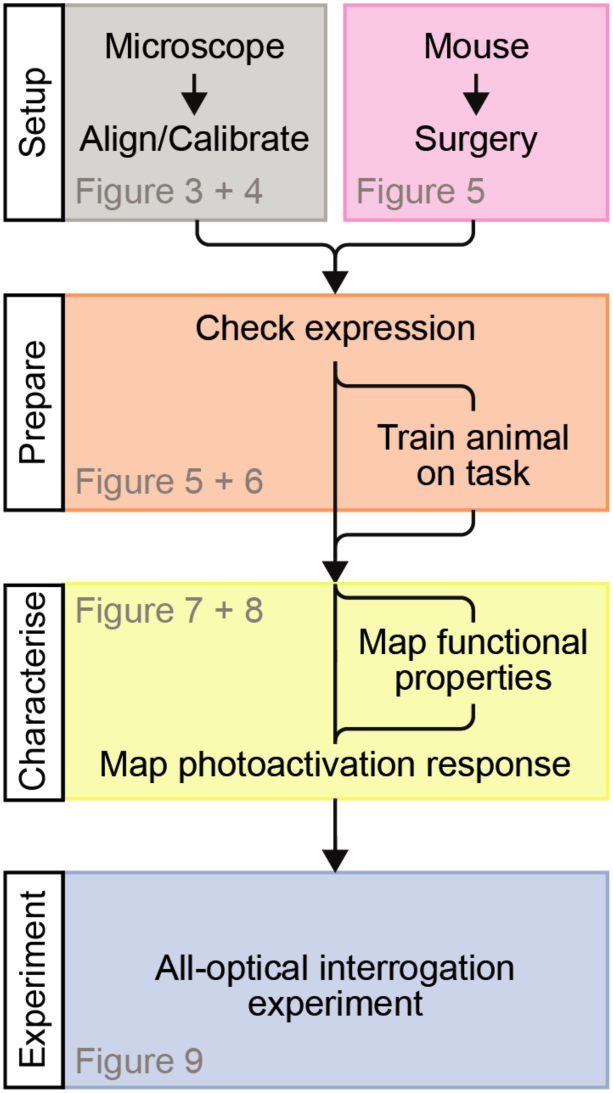
Overview of experimental steps. Essential steps, common to all all-optical experiments. The microscope must be aligned and calibrated before use. Animals used for experiments are engineered to express an activity indicator and opsin in specific neuronal populations. The expression of these constructs is assessed. Animals are (optionally) trained on a behavioural task. Neural responses to a stimulus / task variable of interest are mapped. Neural responses to photostimulation are mapped in order to identify photostimulatable cells. Finally, an experiment can be performed whereby functionally characterized neurons are targeted for photostimulation during a behaviour of interest.

**Figure 3.**
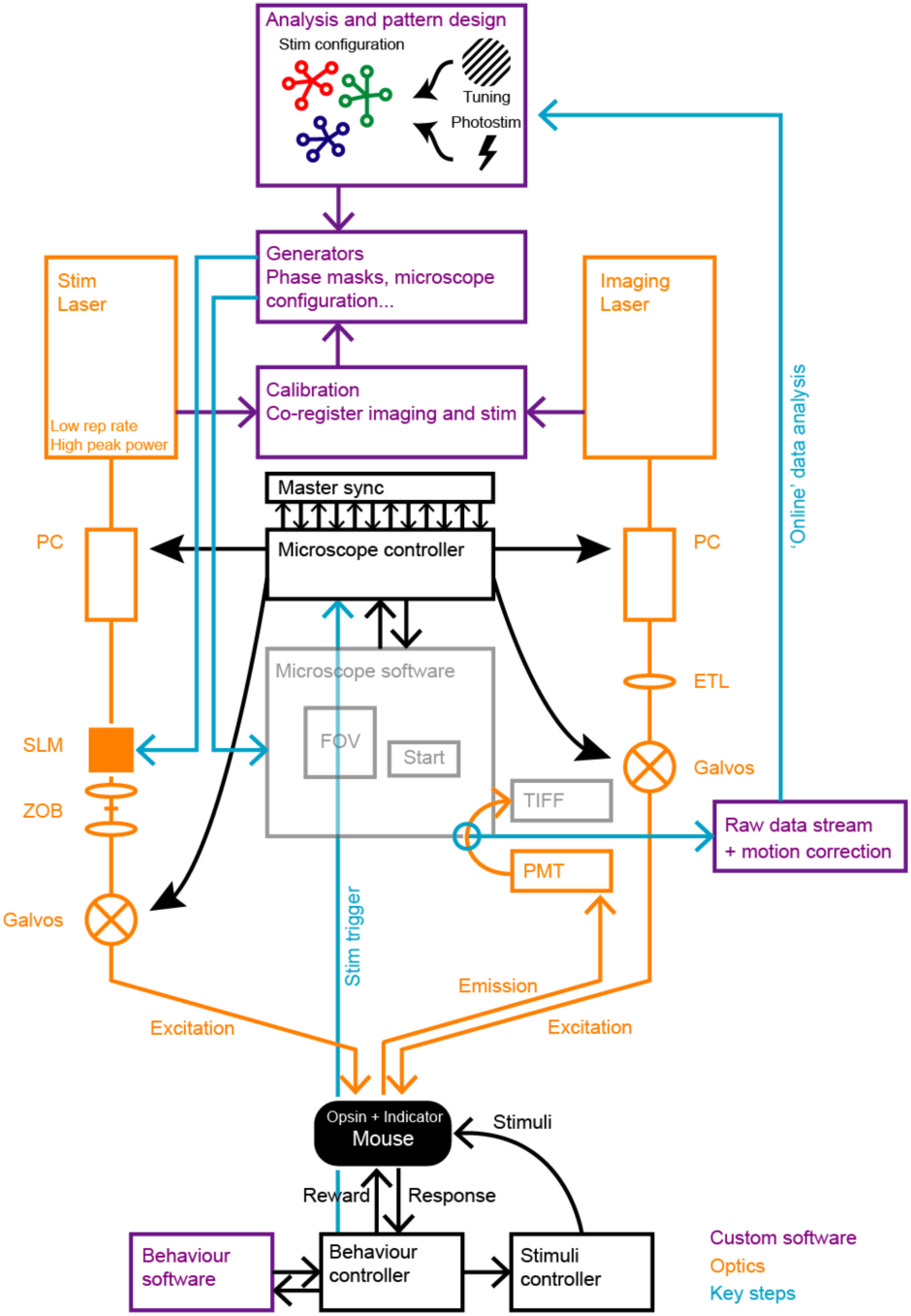
System diagram. To perform all-optical experiments custom software is used to generate stimulation patterns targeted to neurons of interest as identified by analysis of imaging data. The stimulation patterns are generated in the form of files to load into different microscope software modules interfacing with the optical components and requires the use of predetermined calibrations. The stimulation pattern files configure the system such that external triggers, e.g. from a behavioural experiment, can trigger the stimulation of particular neurons by determining the diffraction pattern caused by the SLM and driving power modulation devices as well as galvanometer mirrors.

In this protocol we present a series of steps to enable successful execution of all-optical experiments (**Figure 2**), drawing on our experience with such experiments in five different circuits (L2/3 S1, L2/3 V1, L5 V1, CA1 and CA3). These steps range from the setup and calibration of an all-optical system (**Figure 3, 4**), to the preparation of an indicator and opsin-expressing (**Figure 5**) and task-performing (**Figure 6**) animal, to the characterization of functional (**Figure 7**) and photostimulation (**Figure 8**) responses in a field-of-view (FOV) and concluding with the design and implementation of an all-optical experiment (**Figure 9**). We present the various optimizations that are required for successful implementation of the strategy, as well as highlighting pitfalls and their potential solutions. We have also developed tools for optimizing the experimental workflow, including a strategy for mapping the photostimulation responses of all neurons in a given FOV (using a software package we call Near-Automatic Photostimulation Response Mapper: Naparm), as well as various other software tools used to control our behavioural and all-optical experiments. This protocol (the expression strategies, calibration routines and software and hardware tools described herein) has been instrumental in enabling all-optical experiments in our lab. In combination with the tools developed by other groups, the protocol can form the basis for a standardized toolkit to facilitate the dissemination of these techniques.

**Figure 4.**
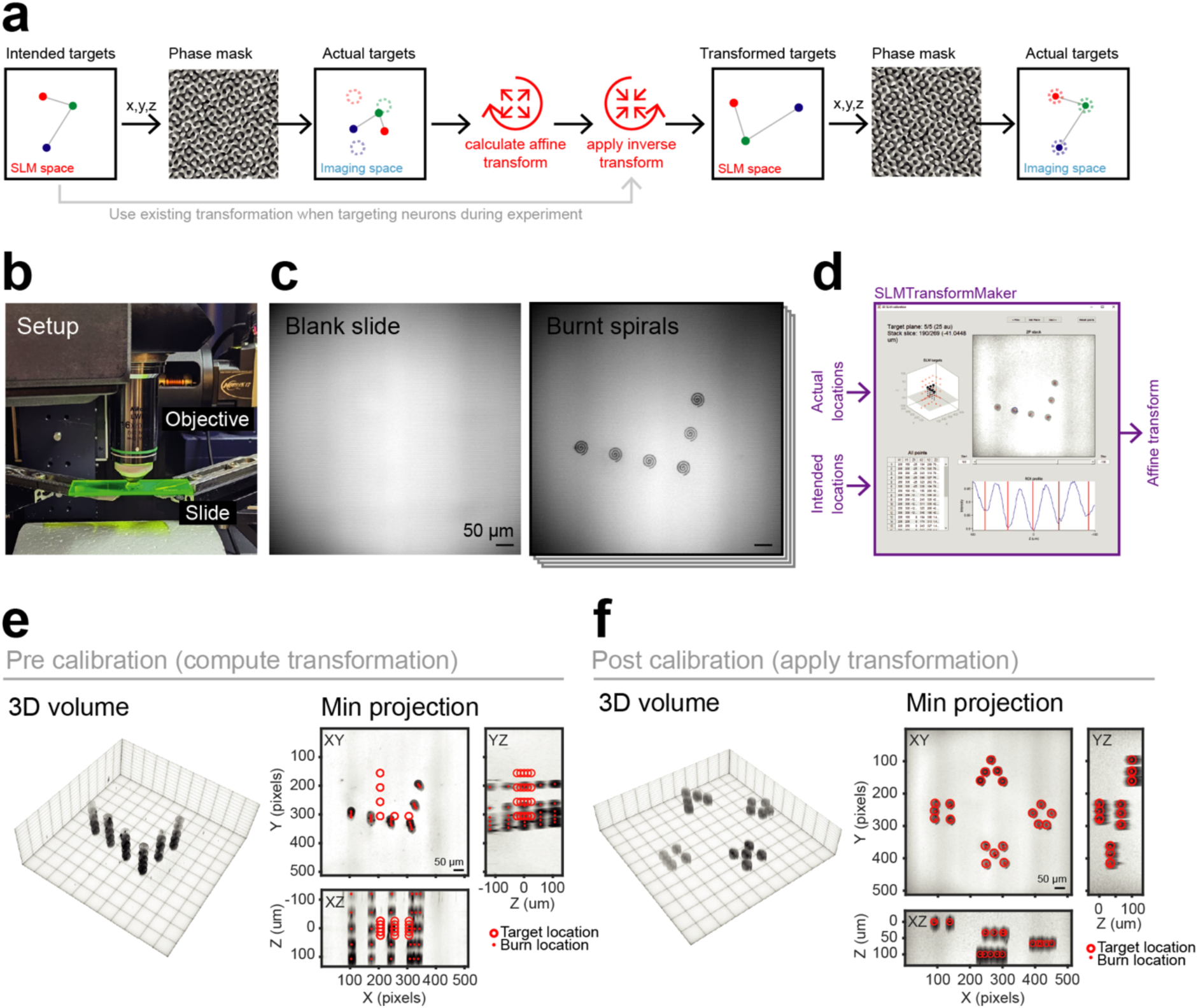
SLM calibration: Mapping photostimulation targets to imaging coordinates. A) Without calibrating the SLM coordinates, the diffraction pattern generated by the SLM will focus in arbitrary locations of the imaging FOV, precluding accurate targeting of precise neurons. By mapping the transformation between programmed SLM coordinates and ultimate location on the imaging FOV, the inverse can be applied allowing for precise targeting of neurons. B) Photograph of fluorescent plastic slide, used for calibration, imaged by the microscope objective. C) Left: image of the fluorescent slide acquired by the imaging pathway. Right: image of the same slide after programming the SLM and burning the resulting spots into the slide by spiral scanning. D) Software used to compute the transformation between SLM target coordinates and the imaging FOV locations of those SLM spots after burning them into a plastic slide. E) 3D projection of a volumetric stack taken of the burnt SLM spots (burnt on 5 axial planes) acquired for the calibration process. F) 3D projection of a different set of SLM patterns, but after calibration, demonstrating successful targeting to intended locations.

**Figure 5.**
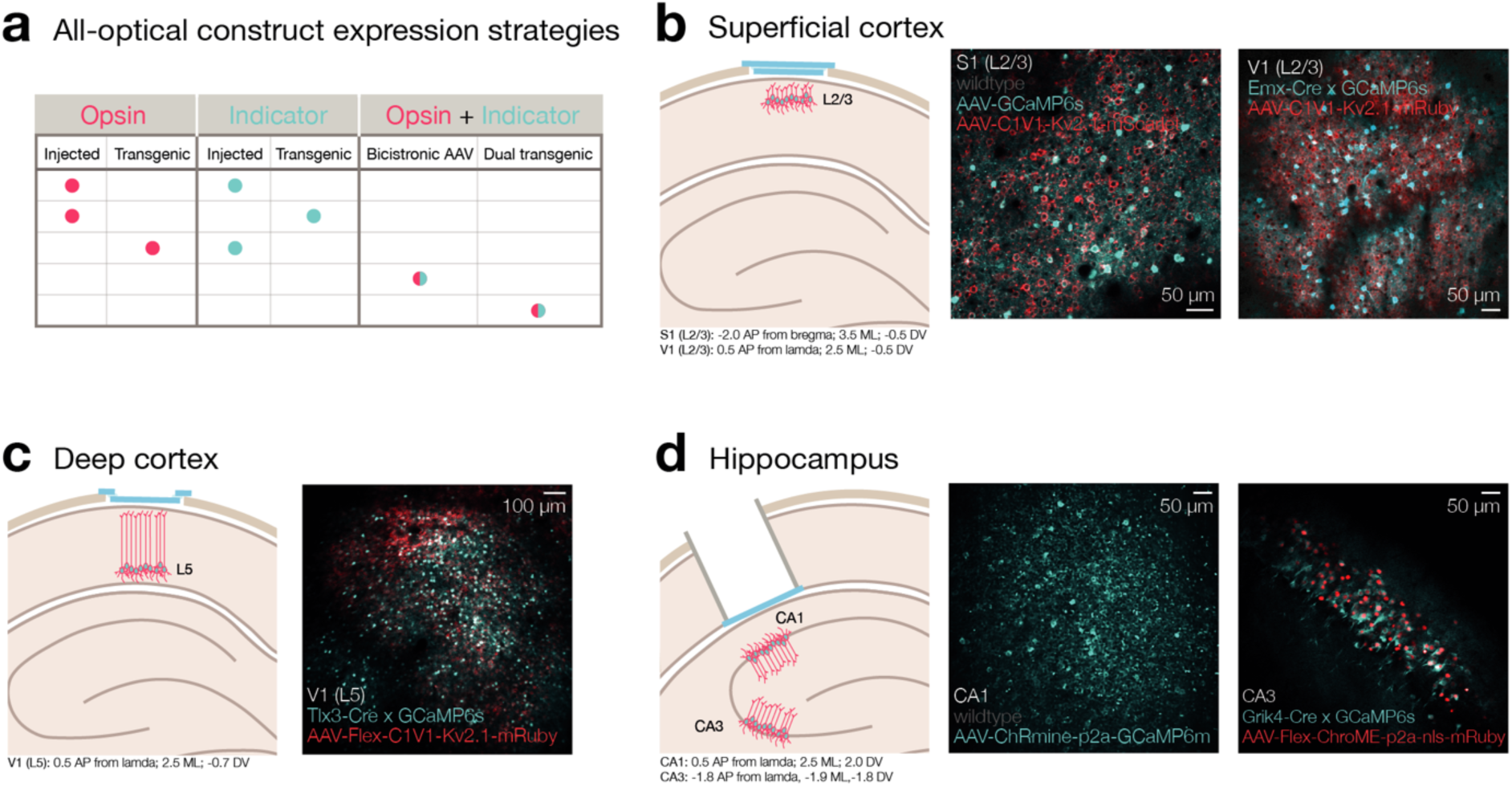
Inducing and checking expression of all-optical constructs. A) Strategies for achieving co-expression of all-optical constructs. B) Co-expression of all-optical constructs in superficial cortex (L2/3 S1 and V1). *Left*: experimental prep schematic; chronic imaging window installed on the cortical surface with either dual AAV expression of GCaMP and C1V1 (S1) or AAV expression of C1V1 in GCaMP transgenic mice (V1). *Right*: example healthy co-expression. C) Co-expression of all-optical constructs in deep cortex (L5 V1). *Left*: experimental prep schematic; chronic imaging window installed on the cortical surface of L5-Cre transgenic mice injected with FLEX-C1V1 and FLEX-GCaMP. *Right*: example healthy co-expression. D) Co-expression of all-optical constructs in sub-cortical structures (hippocampal CA1 and CA3). *Left*: experimental prep schematic; cortical aspiration combined with a canula + chronic imaging window in CA1/CA3-Cre transgenic mice injected with FLEX-C1V1 and FLEX-GCaMP. *Right*: example healthy co-expression.

**Figure 6.**
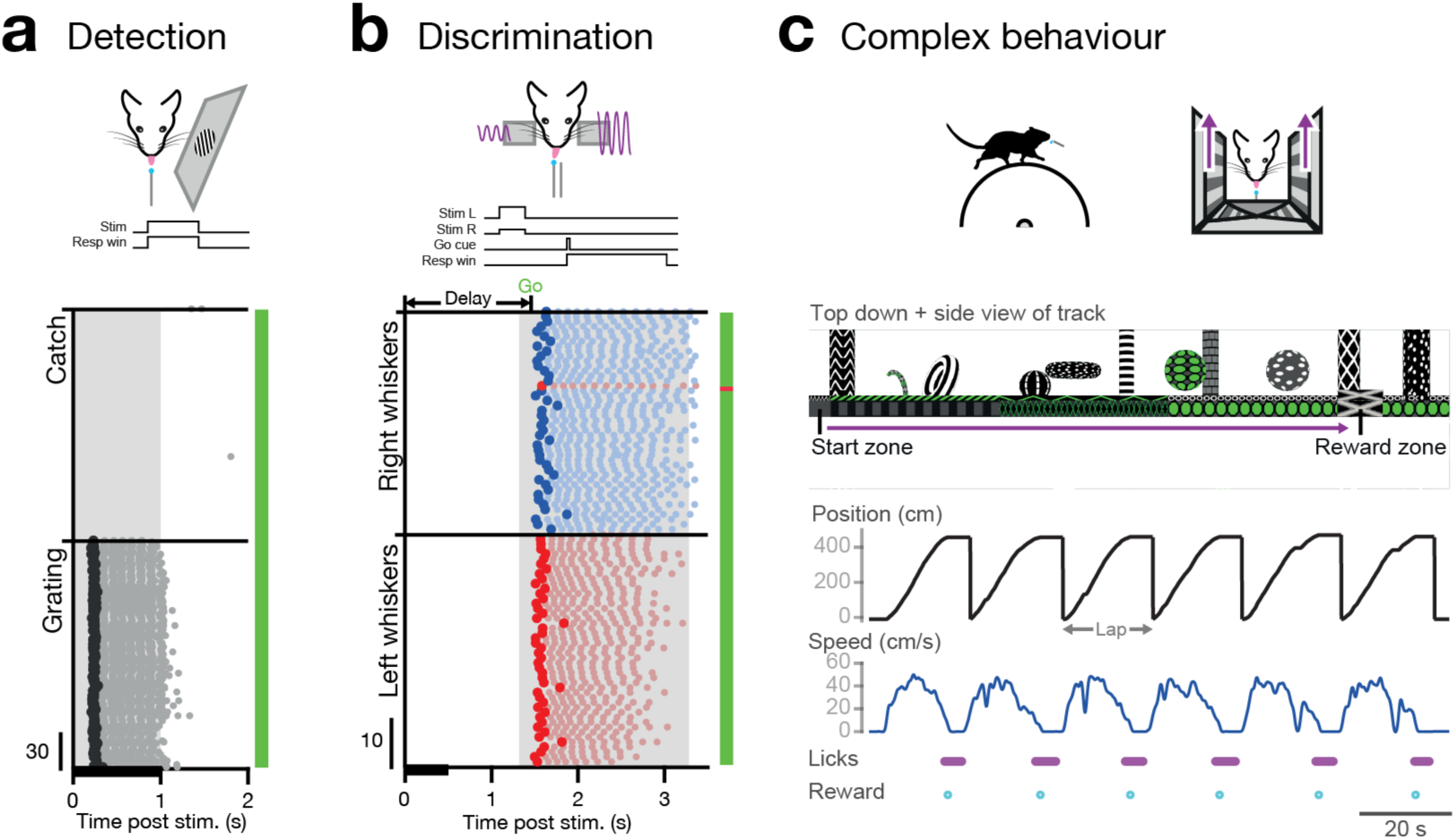
Choosing an appropriate behavioural paradigm. A) Example visual detection behaviour. *Top*: task schematic; mice are required to report the presence of a randomly oriented drifting grating on a monitor by licking for a sucrose reward at an electronic lickometer. *Bottom*: sorted lick raster vertically stacking post-stimulus epochs indicating the stimulus duration (black horizontal bar along x-axis), first lick on each trial (black dot) and subsequent trial licks (grey dots). Task performance is indicated by the colour bar on the right. Note that trials were delivered pseudorandomly but sorted for display. B) Example delay whisker discrimination behaviour. *Top*: task schematic; mice are required to report which whisker pad receives the highest amplitude sinusoidal piezo vibration of two simultaneously delivered bilateral whisker stimuli by licking for sucrose rewards at one of two lickometer ports following an auditory tone Go cue signalling the end of a 1.5 s delay period post-stimulus onset. *Bottom*: sorted lick raster vertically stacking post-stimulus epochs, conventions same as in (A) except that right and left licks are coloured red and blue respectively. Note that trials were delivered pseudorandomly but sorted for display. C) Example complex spatial navigation paradigm. *Top*: task schematic; mice are head-fixed on a cylindrical treadmill that controls movement through a virtual linear world displayed on three surrounding monitors. They spawn at the start of the virtual track and are required to run the length of the track before stopping and licking in the designated reward zone. *Middle*: virtual linear track indicating task features, start zone and reward zone. *Bottom*: position, speed, lick times and reward deliveries for successive laps along the track.

**Figure 7.**
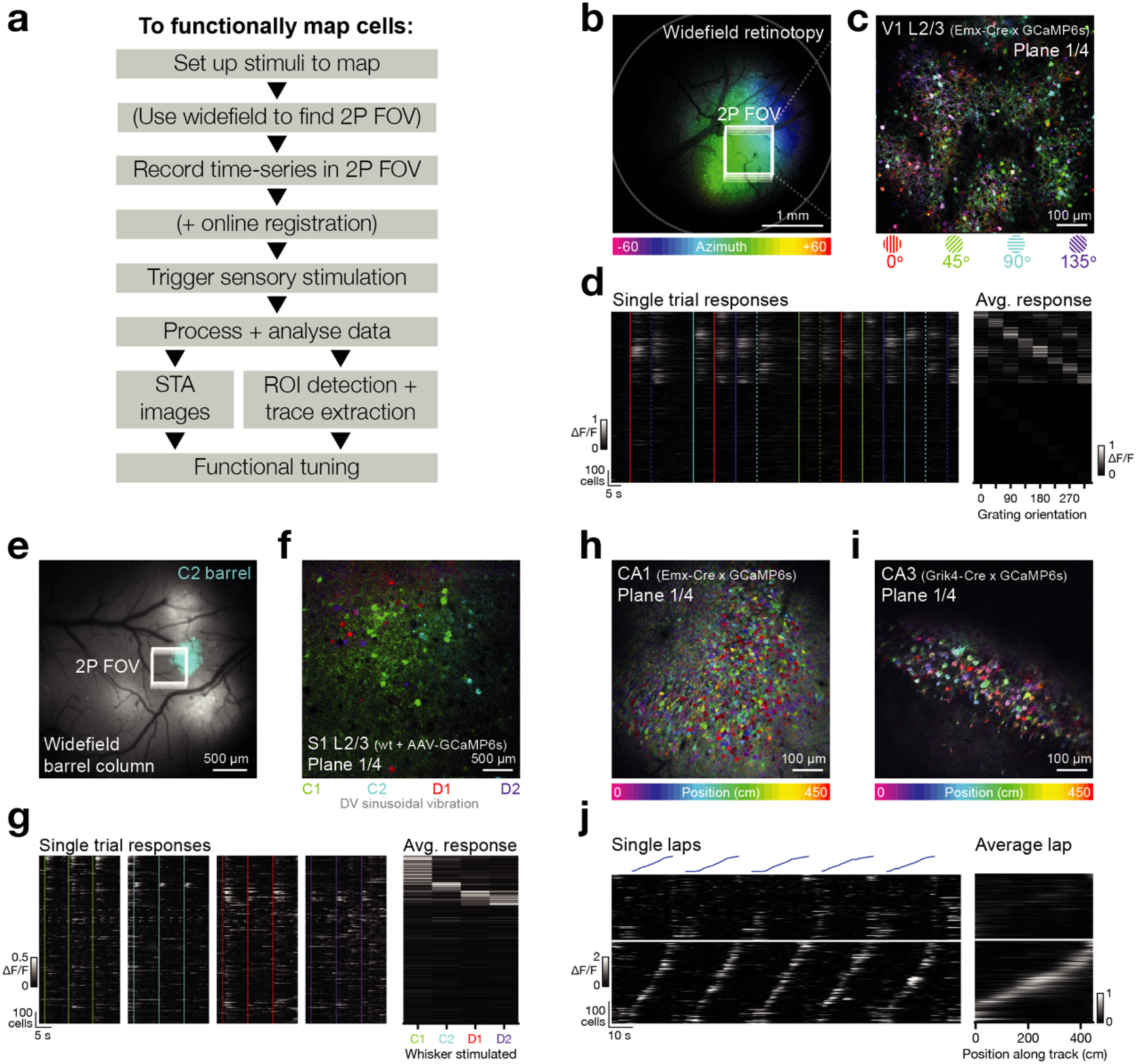
Mapping functional responses online. A) Example workflow for collection and fast online analysis of functional responses. Key optimizations that allow same-session analysis are: (1) online registration in real-time, which eliminate lengthy post-acquisition registration times, and; (2) generation of STA images which intuitively map response strengths and tunings onto the spatial locations of cells. B) STA image generated from widefield calcium imaging data acquired in V1 as a contrast-reversing checkerboard (10°) drifted horizontally across a grey screen (25°/s) positioned in front of the contralateral eye. Pixels are coloured by the azimuth that elicited the strongest response. The 2-photon imaging volumetric FOV used for panels (C) and (D) is indicated. C) STA image generated from one plane in the 2-photon imaging volumetric FOV indicated in (A) (L2/3 V1) as Gabor patches (30°) of drifting sinusoidal gratings (0.04 cycles/°) of 4 orientations (0°, 45°, 90° and 135°) were presented to the contralateral eye. Pixels are coloured by the orientation that elicited the strongest response. D) *Left*: extracted traces showing single trial responses to stimuli indicated by vertical coloured lines (colour conventions same as (C) dashed lines are stimuli in the opposite direction to the solid lines). *Right*: average post-stimulus response amplitude. Note in both heatmaps neurons have been sorted by preferred stimuli. E) Indicator expression image in S1 (greyscale) overlaid with thresholded STA image heatmap (cyan) acquired during vibration of the C2 whisker. The 2-photon imaging volumetric FOV used for panels (F) and (G) is indicated. F) STA image generated from one plane in the 2-photon imaging volumetric FOV indicated in (E) (L2/3 S1) as each of 4 whiskers were stimulated individually (C1, C2, D2, D1). Note this is a composite image combining data from 4 separate movies, one for each whisker. G) *Left*: extracted traces showing single trial responses to stimuli indicated by vertical coloured lines (colour conventions same as (F)). *Right*: average post-stimulus response amplitude. Note in both heatmaps neurons have been sorted by preferred stimuli. H) STA image generated from one plane in the 2-photon imaging volumetric FOV in CA1 (animal genotype: Emx-Cre x CaMKII-tTa x GCaMP6s). Data acquired as animals ran along a virtual linear track. Colour indicates the position along the virtual track that elicited the strongest response, and intensity indicated the response magnitude. I) Same as (H) but in a CA3 FOV (animal genotype: Grik4-Cre x CaMKII-tTa x GCamP6s). J) *Left*: extracted traces showing single trial responses (bottom heatmaps) as an animal ran laps along the virtual linear track (top trajectories). *Right*: response of all neurons averaged across laps. Note in both heatmaps neurons have been first divided into those that are spatially modulated (bottom) and those that are not (top), and sorted within those pools by preferred firing location (place field).

**Figure 8.**
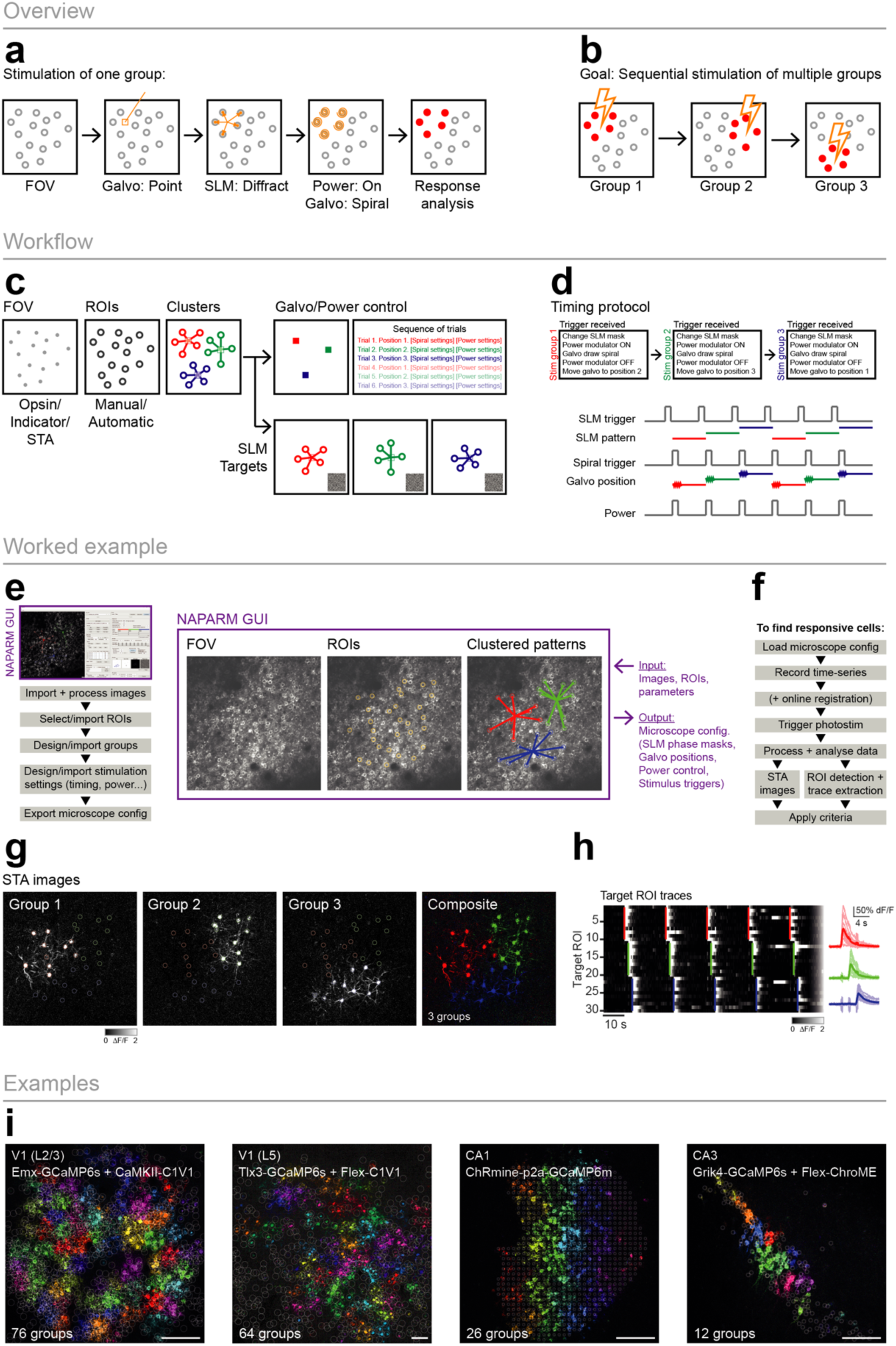
Mapping photoactivatable neurons. A) To stimulate one group of cells the galvanometer mirrors first move to the centroid of the target neurons. The SLM displays a phase mask resulting in diffraction of the beam focusing onto the cells of interest. To stimulate the power modulation device is turned of and the galvanometer mirrors simultaneously move all the diffracted spots in a spiral over the cell bodies of interest. Following stimulation the response can be analysed. B) To stimulate all cells in the FOV sequential stimulation of smaller groups is required. C) The workflow. First, a FOV is loaded and analysed, ROIs are found/selected and then clustered into groups which will be stimulated one after the other. To drive the microscope system to perform the stimulation as described in a) various files are required to configure the subsystems including the positioning of the galvanometer mirrors, the SLM phase masks and a trial sequence listing the stimulation order. D) A voltage command is sent from external hardware to trigger the delivery of a photostimulation. The trigger will update the SLM phase mask, move the galvanometer into position, turn on the power modulation and start the galvanometer spiral. E) Software used to design photostimulation mapping experiments. Workflow follows as: loading FOV images, selecting ROIs, designing the grouping, Configuring the stimulation parameters and finally exporting the files to load into microscope control systems. F) Protocol to run the photostimulation mapping experiment: Load the generated microscope configuration files, record a time-series movie (optionally performing online motion correction), trigger the stimulations throughout the recording. After acquisition the data is analysed in order to identify responsive cells. G) Example STA images of the response following photostimulation of 3 groups (stimulated 1 second after each other). Right shows a composite image where the hue corresponds to pattern number and intensity corresponds to the response magnitude. H) Activity traces extracted from ROIs targeted in the same experiment as in G), stimulations are indicated by vertical coloured lines extending through the neurons that were targeted in a particular pattern. Right shows stimulus triggered average traces. I) Example STA images for the response to photostimulation mapping experiments as in G) in various brain regions.

**Figure 9.**
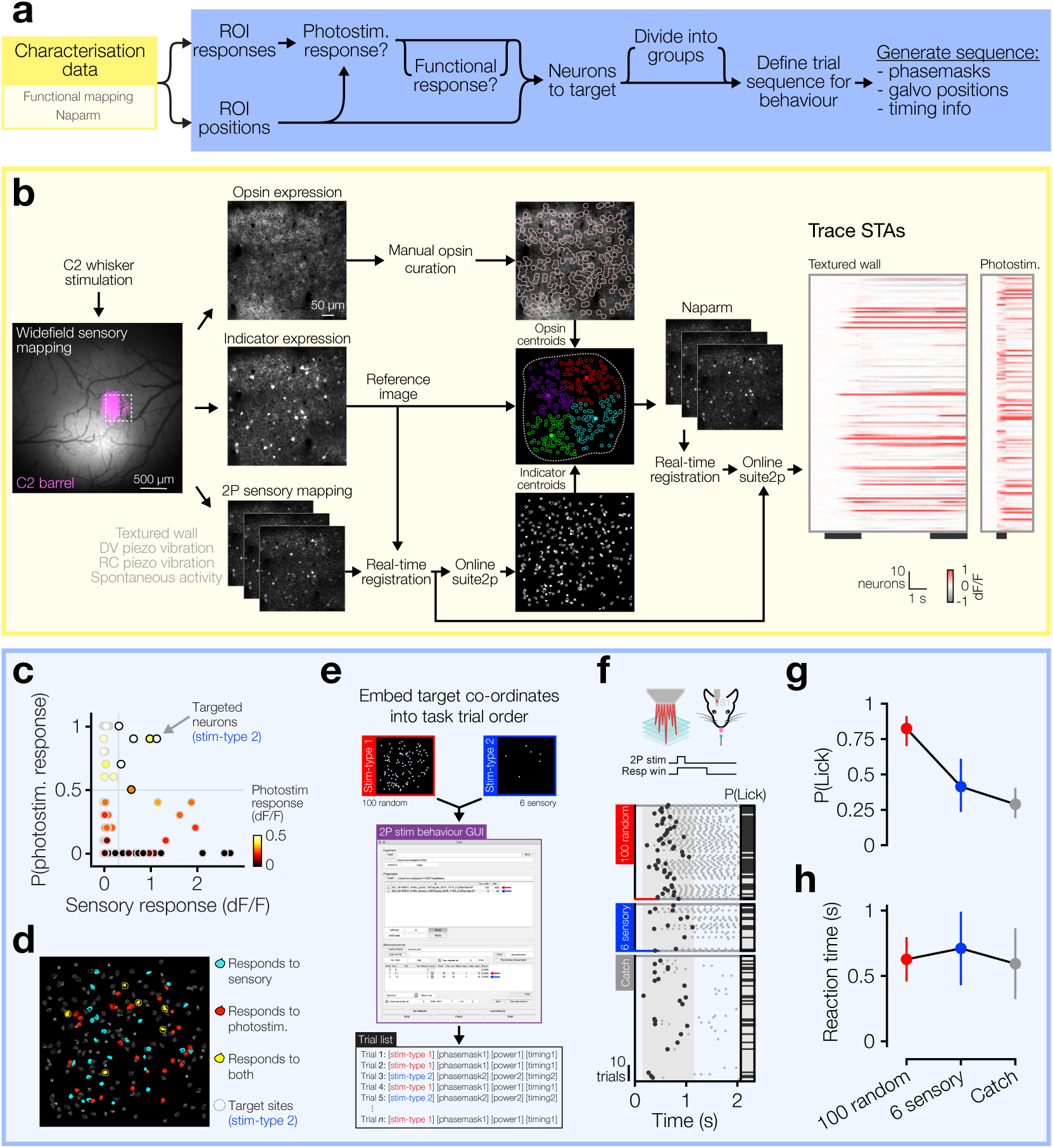
A worked example: probing the perceptual salience of sensory responsive neurons in L2/3 barrel cortex using targeted two-photon optogenetic stimulation. A) Schematic illustrating the general workflow from acquisition of characterisation data to generation of components necessary to perform an all-optical experiment. B) Sequence of steps necessary to acquire relevant characterisation data in this worked example. *Left*: C2 whisker stimulation during widefield calcium imaging of S1 allows identification of the C2 whisker barrel used as the 2-photon FOV going forwards. *Middle-left:* 2-photon expression images of opsin (top), indicator (middle) and 2-photon imaging movies acquired during whisker stimulation (bottom) are used for *Middle-right*: selection of opsin expressing ROIs (top), as a reference image for real-time registration (middle) and to generate functional GCaMP- expressing ROIs via online Suite2p (bottom) with opsin and Suite2p centroids then used to generate 2-photon photostimulation targets (middle). Only targets in the central region of the FOV are included (dashed white border) and these are divided into 4 groups of 50 neurons (colours). These groups are then used for the Naparm protocol which is subsequently concatenated with previously acquired sensory characterisation movies and run through online suite2p. *Right*: This yields extracted traces from Suite2p ROIs which can be used to generate sensory and photostimulus STAs. C) Thresholds (grey dashed lines) are set on the sensory and photostimulus responses to find photostimulable neurons that also respond to sensory stimuli. D) Overlay of sensory and photostimulus response types onto the Suite2p ROI image, highlighting the location of target neurons (dashed circles). E) Target co-ordinates for two photostimulus trial types, one stimulating 100 random neurons (trial-type 1) and another stimulating just the 6 sensory responsive neurons (trial-type 2; see C – D), are embedded into a behavioural task paradigm via a custom GUI that allows the binding of specific phase masks with trial types and the organisation of trial types into a sequence of trials for a given behavioural session. F) *Top*: task schematic; mice are required to report the detection of 2-photon photostimulation targeted to ensembles of neurons (500 ms duration; 10 x 20 ms spirals at 20 Hz) by licking at an electronic lickometer for sucrose rewards in a 1 s response window following the onset of photostimulation, or to withhold licking on catch trials during which no neurons were photostimulated. *Bottom*: sorted lick raster split by trial-type from the behavioural session immediately following the characterisation in B – E. Trials were delivered pseudo-randomly but are sorted for display. Stimulus durations shown as coloured bars at bottom of raster. G) Proportion of trials on which the animal licked, and therefore putatively detected photostimulation, for each trial-type. Error bars are binomial. H) Reaction time for each trial-type. Error bars are s.d.

### Experimental design

#### System design

The ‘all-optical’ method combines genetic engineering of neurons, multiphoton imaging and optogenetic manipulation, and holographic optics for optical recording and targeted photostimulation. Integrated hardware and software is needed to coordinate all aspects of the system in order to read and write neural activity in awake, behaving, animals. In this section we will give an overview of the system design and then provide more detail in subsequent sections.

To enable light-based readout and control, neurons need to be engineered to co-express two proteins: a fluorescent indicator to record their activity, and an opsin to manipulate their activity. These two proteins should be spectrally separated, meaning a different wavelength of light is used to excite the activity-dependent fluorophore than is used to activate the opsin. Two-photon excitation of both the indicator and the actuator is used to provide good depth penetration and optical sectioning for both the reading and writing channels.

A core element of the approach is that two-photon imaging of large populations of neurons is performed simultaneously with two-photon stimulation of those or other neurons. This simultaneity is made possible by two independent lightpaths which are combined through the same objective. Firstly, an imaging lightpath that resembles a conventional two-photon scanning microscope. And secondly, an additional lightpath devoted to photostimulation, requiring an additional laser source specifically suited for optogenetic stimulation, a power modulation device (a Pockels cell or an acousto-optic modulator (AOM)), a programmable diffractive element, typically a spatial light modulator (SLM), that allows the targeting of multiple neurons, and (optionally) a set of galvanometer mirrors to ‘spiral’ the focused points of light over neuronal cell bodies to excite sufficient numbers of actuator (opsin) molecules (J. P. Rickgauer and Tank, 2009). These two lightpaths must employ separate lasers of sufficiently different wavelength (in part dictated by the choice of indicator and opsin) in order to avoid ‘crosstalk’ – the unintentional activation of opsin-expressing neurons by the imaging laser. The optical design of the stimulation path typically mirrors that of a conventional imaging pathway, with the addition of the SLM at a plane conjugate to the galvanometer mirrors and the objective back aperture. SLMs enable user-programmed diffraction of the stimulation beam into multiple beamlets targeted to many individual neurons at once. This diffractive action is not 100% efficient: the light that is not diffracted remains in the 0^th^ diffraction order (‘zero order’) and needs to be blocked before entry to the objective to prevent nonspecific stimulation of neurons. To block this ‘zero order’ we position a small physical blocking element (as the incident power can be high and is localised in a focal point we use reflective materials such as a piece of aluminium foil, or lithographically printed gold deposits, rather than absorbent materials) installed in a translatable mount at the focal plane of the SLM, preventing the ‘zero order’ from propagating further through the rest of the system. While we, and most commercial systems, use an SLM in combination with galvanometer spiral scanning to rapidly illuminate multiple cell bodies, this approach is not the only solution. Galvanometer mirrors, without an SLM, can be used to sequentially target single cells one at a time (Rickgauer *et al*., 2014; Chettih and Harvey, 2019; Jennings *et al*., 2019). Additionally, a diffraction grating, with or without an SLM, can be used to generate a temporally-focused, arbitrarily shaped blob of light which illuminates laterally extended areas (e.g. the whole soma) at once, negating the need to ‘spiral’ the beamspots (Papagiakoumou *et al*., 2008, 2020; Rickgauer *et al*., 2014; Pégard *et al*., 2017) (**see Box 1**).

To perform an all-optical experiment involving targeting of functionally identified neurons, online analysis is required to identify, characterize and then stimulate the cells of interest or relevance (**Figure 3**). For many experiments, we rapidly perform the analysis immediately after the acquisition of data has completed while the animal remains mounted under the microscope. The speed of this analysis is vital to maintain performance of animals in behavioural experiments and is facilitated by real-time access to the incoming data stream; the raw imaging data is streamed to an immediately readable raw binary data file (avoiding slow TIFF read and write overheads) while also performing real-time motion correction frame by frame as the data is being acquired. These optimizations allow for analysis of the neural data immediately after acquisition.

To identify recorded neurons and extract their activity through time we either use a version of Suite2p (Pachitariu *et al*., 2016) modified to work on the real-time registered binary files mentioned above, or semi-automatically select regions-of-interest (ROIs) from pixelwise stimulus triggered average (STA) images of the FOV responses to stimuli of interest. With access to the raw data stream it is also possible to algorithmically identify ROIs as the data is being acquired (Giovannucci *et al*., 2019) and/or readout and manipulate activity in closed-loop (Grosenick *et al*., 2015; Zhang *et al*., 2018) allowing the real-time neural activity, or animal behaviour, to guide the optogenetic manipulation. Finally, deciding on the pattern of target neurons to stimulate depends on three factors: physical location, functional identity (i.e. tuning to a task/stimulus variable of interest), and whether they are responsive to photostimulation. Given these parameters, we construct a target pattern and generate requisite files to be loaded into the respective modules of the all-optical system which include: the diffraction pattern to be made by SLM, the instructions for the microscope software to position photostimulation galvanometer mirrors and control the photostimulation power modulator, as well as triggers to synchronise the pieces of hardware. To enable subsequent analysis we typically record signals of all imaging frame acquisitions, stimulation triggers and behavioural events by one master ‘synchroniser’ data acquisition (DAQ) device.

#### Choice of indicator/opsin combination

All-optical constructs should be carefully chosen and evaluated to balance the sensitivity of the indicator for reporting spikes and the efficacy of the opsin for two-photon activation whilst ensuring the maximal spectral distance between the two excitation wavelengths to minimise ‘crosstalk’ – the unintentional activation of opsin-expressing cells by the laser used to image the activity indicator. For this reason, green indicators (which are excited by blue one-photon light), such as those of the GCaMP family (Chen *et al*., 2013) (optimal two-photon λ_excitation_ ∼920 nm), should generally be used with red-shifted opsins such as C1V1 (Yizhar, Lief E. Fenno, *et al*., 2011), Chrimson (Klapoetke *et al*., 2014), or ChRmine (Marshel *et al*., 2019) (two-photon λ_excitation_ ∼1000-1100 nm). Opsins sensitive to blue light with very fast channel kinetics, such as ChroME (Mardinly *et al*., 2018), can also be used if imaging power and dwell time are sufficiently low (Mardinly *et al*., 2018). Red indicators (which are excited by yellow one-photon light), such as jRCaMP, or jRGECO (Dana *et al*., 2016) or xCaMP (Inoue *et al*., 2019), should generally be used with blue-light sensitive opsins (Forli *et al*., 2018, 2021) such as ChR2 or ChroME. Different experiments may require opsins with different features: for experiments where spike timing with millisecond precision or specific spiking rate are important variables then opsins with fast off channel kinetics (on the order of 5-10 ms) are required to ensure high temporal fidelity (such as Chronos or ChroME), whereas if only the number or identity of cells is important, with little emphasis on temporal features of their activity, opsins with slower temporal characteristics (e.g. C1V1) can be used. It is important to note that there is typically a trade-off between opsin speed and sensitivity. Since fast opsins will not integrate current over time they must be sensitive and/or have high conductance to allow for sufficient depolarisation during the short, temporally precise stimulation times that they allow. Additionally, in this protocol we provide examples where we have extended the genetic engineering approach to use the Cre/LOX system to restrict expression to specific cell-types of interest.

The opsin molecules are typically coexpressed with a fluorescent marker in order to visualise expression. These markers can be directly fused to the opsin in the membrane (e.g. C1V1-Kv2.1-mRuby) or they can be separate from the opsin molecule through the use of self-cleaving peptide links (e.g. C1V1-p2a-mCherry) resulting in the opsin remaining in the membrane, but the fluorophore free in the cytosol. The reporter fluorophore can be restricted to the nucleus which can help in cell identification (e.g. ChroMe-p2a-nls-mRuby). Finally, the opsin can be expressed from a bicistronic construct also containing the indicator gene and thus require no additional fluorophore for identification (e.g. GCaMP6m-p2a-ChrMine).

A major determinant of the fidelity, or spatial resolution, of photostimulation is the localization of opsin expression. If opsin is expressed in the membranes of all neuronal processes, photostimulation directed to a location far from a given cell body may still depolarize that neuron if it has processes passing through the stimulation volume. To this end, multiple groups have developed somatic restriction strategies (Lim *et al*., 2000; Baker *et al*., 2016; Shemesh *et al*., 2017; Mardinly *et al*., 2018; Chettih and Harvey, 2019; Marshel *et al*., 2019), whereby opsin molecules are localized to and concentrated in the soma and proximal dendrites, rather than throughout the dendritic arbor.

The optogenetic toolbox is continually expanding which precludes including a definitive list here. Fortunately, concerted efforts by many groups are constantly yielding up-to-date, useful resources detailing opsin spectral responsivity (relevant to laser choice and crosstalk), sensitivity (relevant for degree of activation and crosstalk) and kinetics (relevant for timing, photostimulation strategy and crosstalk) (Yizhar, Lief E Fenno, *et al*., 2011; Mattis *et al*., 2012; Prakash *et al*., 2012; Wiegert *et al*., 2017; Antinucci *et al*., 2020; Sridharan *et al*., 2021) that can be used for making informed decisions about which construct is appropriate for a setup.

After having chosen an opsin and indicator pair, the next most important step is to optimize the level of expression. Achieving balanced co-expression is challenging, perhaps due to promotor-specific differences in different brain areas, but also likely due to overexpression of one or both constructs (Tian *et al*., 2009; Chen *et al*., 2013). Due to factors like competition between different viruses, and depending on the experimental goal of long-term health versus maximum possible expression, all combinations of chosen indicator and opsin must be thoroughly tested in the particular preparation of interest at a range of titres over different timescales in order to assess efficacy (avoiding under-expression) and long-term health (preventing over-expression (Tian *et al*., 2009; Chen *et al*., 2013): see also Supp Figure 8 in Packer et al. 2015).

#### Choice of photostimulation laser

It is important to choose the photostimulation laser carefully to optimize the activation of opsin-expressing neurons. The two-photon absorption by opsins (and thus activation of neurons) is proportional to the square of the excitation light intensity (**Box 2**), and the most optimal stimulation will be achieved with wavelengths closer to the peak of their excitation spectrum. Pulsed lasers for two-photon photostimulation are described by a few key parameters, namely: wavelength, peak power per pulse, pulse repetition rate and pulse width (which together dictate the average power). We discuss these parameters in detail below.

Laser wavelength for photostimulation (either when selecting a fixed-wavelength laser, or when using a tunable laser) should be chosen with respect to the peak of the absorption spectrum of the opsin of choice. Although in practice, given the prevalence of low rep rate lasers in the >1000 nm range, opsins are often chosen to match available lasers. The laser power at the wavelength closest to the peak of the opsin absorption spectrum will dictate the number of neurons that can be activated simultaneously, since the total power will be divided (usually equally) amongst the holographically split beamlets targeted to individual cells.

Pulsed lasers used for two-photon imaging typically operate at a high repetition rate (∼80 MHz), which is necessary given the speed at which the focused beam is scanned across the tissue of interest dictating the ‘dwell time’ for fluorophore exposure. This high repetition pulse rate is associated with a trade-off between the peak power – power delivered by each pulse – and the time-averaged power (see **Box 2**). While similar lasers (Fianium [2 W average power, 80 MHz rep-rate], Coherent Fidelity [2 W average power, 80 MHz rep-rate]) have been used for photostimulation of opsin expressing neurons, we and other groups have found more success with a different type of laser. Low-repetition rate lasers (e.g. Amplitude Satsuma [1060 nm, 20 W average power, 0.5 MHz/2 MHz rep-rate]; see also Coherent Monaco, Menlo BlueCut, SpectraPhysics Spirit and alternatives) are associated with a much larger peak power (compared to high-repetition rate lasers) while maintaining a similar average power. High peak powers more efficiently activate opsin molecules by means of two-photon absorption while opsin and cellular integration kinetics allow for the reduced frequency of pulses. Using pulses with higher peak powers means less average power is required to successfully stimulate a cell. Less average power translates to less thermal energy and thus less heating of the tissue (which could lead to thermal damage such as protein denaturation) (Podgorski and Ranganathan, 2016; Picot *et al*., 2018). At a given average power, higher peak powers enable the beam to be split across more neurons to be stimulated simultaneously. Note however that high peak powers can be associated with non-linear damage mechanisms (Koester *et al*., 1999; Hopt and Neher, 2001). Our average power per cell (excitatory cortical L2/3 cells expressing C1V1 opsin) of ∼6 mW at a repetition rate of 2 MHz was selected as a compromise between good photostimulation efficiency and minimal photodamage in our experimental configuration (see **Box 2**). This compromise depends on several parameters, including the sensitivity of the opsin and the imaging depth in tissue and should be tested for each setup (for example, for our deeper cortical L5 experiments we typically use ∼12 mW per cell at 1 MHz repetition rate).

#### Patterned illumination device

We use a reflective phase-only LCoS (liquid crystal on silicon) SLM to introduce diffraction patterns onto the photostimulation laser beam which is focused into spots of light on the sample targeted to specific neurons. We then use a pair of galvanometer mirrors to scan these spots of light over the cell bodies of these neurons in a spiral pattern to illuminate as many cell-membrane localised opsin molecules as possible.

Reflective phase-only SLMs are able to modify the phase of the wavefront reflected off them. This is achieved through the action of birefringent liquid crystals in the active surface of the SLM. The SLM active surface is composed of an array of pixels, where each pixel is an electrode that controls the orientation of the liquid crystals above it. Depending on the orientation of the crystals (controlled by the voltage applied to the electrode) the optical path length is increased or decreased resulting in the light passing through those crystals having a modified phase relative to the incident beam. This phase modulated wavefront is Fourier transformed by the objective into multiple foci in the sample. The overall effect of the SLM on the laser wavefront is governed by the coordinated action of all its pixels, which are controlled through addressing the SLM with a ‘phase mask’ (also referred to as a hologram), which in effect controls the voltages applied to each pixel electrode. These phase masks can mimic the action of physical optical elements such as lenses to focus light or diffraction gratings to diffract light at a particular angle. In practice these customisable phase masks are typically generated with a variant of the Gerchberg-Saxton algorithm (Gerchberg and Saxton, 1972), an iterative Fourier, inverse-Fourier transform procedure (computation of which can be performed on GPUs), but other methods are available (Eybposh *et al*., 2020).

There are a number of operating characteristics to consider when selecting an SLM:

1. Overall efficiency: SLMs are not 100% efficient devices and thus some power is lost by using them. Modern devices are specified as ∼70-90% efficient, though this efficiency is reduced at wider angles of diffraction. The light that is not diffracted remains in what is called the ‘zero-order’ (which is blocked before entry to the objective).
2. The size of the individual pixels: smaller pixels (keeping the SLM size fixed) will allow for greater diffraction angles to be achieved – increasing the size of the addressable FOV under the objective – but may come at the cost of crosstalk between neighbouring pixels, resulting in reduced diffraction efficiency and thus a less efficient hologram.
3. The size of the SLM: this dictates the amount of de/magnification necessary to propagate through the rest of the optical system. The level of magnification will affect the size of the beam at the objective back aperture thus affecting the effective numerical aperture (NA), while at the same time higher magnification will result in smaller diffraction angles being achieved by the objective. There is therefore a trade-off between addressable FOV size and optical resolution.
4. The speed of the SLM: the speed at which the pixels can be driven to a new voltage setting (refresh rate of the driving electronics) as well as adopt a new voltage setting (liquid crystal settling time, i.e. the time for the liquid crystals to reorient to the newly applied voltage) will together dictate the rate at which new diffraction patterns can be focused on the sample, which in combination with exposure durations (see opsin sensitivity) will dictate the speed at which sequences of activity can be ‘played in’.

Alternative systems for patterned illumination can be used. Galvanometer mirrors alone are appropriate if one only needs to stimulate one cell at a time (Rickgauer *et al*., 2014; Chettih and Harvey, 2019; Jennings *et al*., 2019), whereby the focused beam spot can be steered to individual locations in sequence. Digital micromirror devices (DMDs) can also be used to target multiple locations simultaneously (Bhatia *et al*., 2021), as an alternative to SLMs. DMDs can be driven much faster than current SLMs (e.g. > 1 kHz), however, they are less efficient than SLMs as the light that would fall on untargeted regions is discarded rather than refocused to the desired target locations.

#### Choice of two-photon photostimulation method

The high resolution of photostimulation with a two-photon beam spot comes at the price of a very small activation volume (a region of membrane on the order of the point spread function). To increase this volume sufficiently to cause large enough depolarization and therefore generate action potentials one can either use spiral scanning or beam-shaping with temporal focusing (see **Box 1**). We have used spiral scanning (J. P. Rickgauer and Tank, 2009), which involves scanning the beam spot over the somatic membrane, as this strategy requires less average power to cause neurons to spike. Beam-shaping (by under-filling an objective or using digital holography) allows users to increase the lateral extent of the two-photon point spread function (PSF) to be the size of a neuron, but at the cost of also increasing the axial extent (beyond the size of a neuron) which will degrade the resolution of stimulation. Temporal focusing (via a diffraction grating placed in the beampath) can improve the axial extent of the PSF by ensuring the laser pulses have the shortest duration at the focal plane of the objective, with the pulses broadening rapidly along the axial direction. The short pulses at the focus provide greater two-photon absorption relative to those out of focus, confining the excitation volume and recovering stimulation resolution at a cost of having to use more power per cell.

#### Choice of two-photon stimulation parameters

To provide precise control of the photostimulation of neurons, we can adjust the average power per cell, the duration of a single exposure, and the timing of trains of exposures (inter-exposure interval as well as the total duration). For photostimulation response mapping using the relatively slow opsin C1V1, we generally photostimulate with a 20 ms spiral repeated 10 times at 20 Hz, resulting in a 500 ms stimulus epoch. These spirals are approximately a cell diameter in size (on the order of 10 – 15 µm), consist of 3 revolutions and an average power on sample of 6 mW per cell (with a 2 MHz rep rate laser). The stimulation parameters (pulse repetition rate and average power, keeping the stimulus duration constant) were chosen as a result of careful calibration, ensuring efficient activation but also minimising any signs of photodamage (see **Box 2**). In our calibration protocol we varied only one parameter at a time, and started at a lower bound for the value of that parameter. We typically stimulate in a given condition for 10 repeats, then increment the parameter and stimulate 10 more times, repeating until we’ve reached the upper bound of the given parameter. Then, by analysing the resulting data we can arrive at an optimal value for that parameter that gives adequate activation and minimal signs of damage. Our chosen values correspond to a stimulation rate considerably above the median firing rate of neurons in (vS1) cortex while also being within the physiological range of pyramidal cells over short time periods. The stimulation parameters should ideally be assessed for every system and opsin used to ensure effective but safe stimulation. The duration used for a single spiral scan depends on the sensitivity of the opsin used (i.e. the two-photon cross section and the size of the induced photocurrent) and the intensity of light used during exposure. New, more sensitive opsins such as ChRmine (Marshel *et al*., 2019) are compatible with much shorter spirals (sub-millisecond exposures).

Another important characteristic of opsins is their off kinetics. Slower opsins (like C1V1 compared to ChroME) take comparatively longer to close after opening, meaning they are unable to elicit another action potential as quickly. These slower opsins will not be able to faithfully follow high frequency (> 40 Hz) trains of stimulation, though their ability to integrate current over time can mean that they require lower powers to generate sufficient depolarisation if photostimulation duration is not a concern. Additionally, all opsins have the potential to desensitise, that is, become unable to open again after being repeatedly exposed with light of a strong intensity or for a long duration. If the stimulation volume is not saturating (i.e. activating all available opsin molecules) then other opsin molecules can diffuse and replenish the supply mitigating the concern at least partially. The off-kinetics and the rate of desensitization necessitates thought into how frequently neurons are stimulated both within a trial and the time between trials.

As all of these stimulation parameters are under the experimenter’s control the stimulation can be designed to be physiological (replicating naturally occurring firing rates or ensembles of cells) or not, depending on the experimental question.

#### Characterization of the all-optical system

There are two main issues to consider when evaluating the performance of an all-optical system for use in biological experiments. Those are 1) the achievable resolution of photostimulation – i.e. the specificity with which single neurons can be targeted without affecting their neighbours, and 2) the amount of crosstalk between the imaging and photostimulation channels – i.e. the degree to which the laser used to record activity in the FOV also excites the opsin. Both of these considerations depend on both optical and biological factors.

The resolution of photostimulation is determined by the effective NA of the photostimulation light path, which is set by the size of the beam at the back aperture of the objective. The degree to which the photostimulation beam fills the back aperture of the objective represents a trade-off with the maximum achievable diffraction angles, with more magnification resulting in shallower angles. The optical resolution of the system should be assessed empirically by using a sample of small (< 1 µm) fluorescent beads to reconstruct the point spread function following standard procedures. As discussed above a large determinant of the effective resolution is also biological in nature – the expression pattern of the opsin. Neurons are axially extended structures, and therefore if opsin is expressed through the extent of a neuron this can exacerbate poor axial resolution. Restricting the opsin to just the cell body (as discussed above) can mitigate this concern. To accurately quantify the resolution of the system we recommend performing patch-clamp recordings of single neurons to provide electrophysiological ‘ground truth’ for photostimulation (Packer et al. 2015; Mardinly et al. 2018). This involves delivering photostimuli at various lateral and axial offsets with respect to the cell body of the electrophysiologically recorded neuron. The resulting dataset can be used to construct a curve of action potential activation as a function of offset from the soma, and from this curve we can calculate the full width at half maximum (FWHM), a standard measure of the resolution (or point spread function) of optical systems. Using this measurement we can subsequently define regions around the targeted sites in all experiments to either include or exclude neurons from data analysis based on whether they might have been directly stimulated. This is particularly important for studies examining the synaptic recruitment of other, non-targeted neurons in the local neural circuit.

The crosstalk between the imaging and stimulation pathways is largely set by the excitation spectra and sensitivity of the opsin molecule. If the opsin used is highly sensitive and/or has a spectra that overlaps with the imaging wavelength, the imaging laser may activate the opsin and thus change the resting potential. Care needs to be taken to minimise the imaging laser exposure time or exposure intensity, for example by using the minimum imaging laser power that results in usable data. Additionally by scanning volumetrically (in 3D) we can reduce the time the imaging laser is focused on any given cell, thereby reducing the time the opsin molecule are potentially excited. To accurately quantify the degree of crosstalk in a new system we again recommend performing electrophysiological recordings of an opsin expressing cell. By imaging the FOV at increasing imaging laser powers while recording the opsin-expressing cell it is possible to assess how much the resting potential or firing rate changes as a function of the imaging laser. By testing various configurations (scanning speed, FOV size, volumetric scanning, illumination power) it is possible to design an imaging condition that has minimal impact on the baseline properties of opsin-expressing neurons

#### Choice of brain region

The all-optical protocol reported here can in principle be performed in any structure that is accessible to two-photon microscopy. We have carried out successful experiments in L2/3 vS1, L2/3 V1, L5 V1, as well as hippocampal regions CA1 and CA3 (through an implanted cannula). All-optical interrogation of neural circuits has also been applied in the olfactory bulb (Gill *et al*., 2020), orbitofrontal cortex (OFC) (through a GRIN lens) (Jennings *et al*., 2019) and anterior lateral motor cortex (ALM) (Daie *et al*., 2021). The main region-specific considerations are that deeper areas will require higher laser power for both imaging and photostimulation, and that the genetic identity of cells in different regions might preclude the use of some promoters for indicator/opsin expression. Areas deeper than ∼500 µm are typically accessed through either an implanted cannula (Dombeck *et al*., 2010) or using GRIN lenses (Levene *et al*., 2004; Jennings *et al*., 2019), or with three-photon imaging (Horton *et al*., 2013; Ouzounov *et al*., 2017; Wang *et al*., 2018; Weisenburger *et al*., 2019).

##### Box 1 Photostimulation Strategies – Spots or Blobs

Excitation strategies used for two-photon optogenetic stimulation of individual neurons fall into two broad categories: (1) **serial** excitation which relies on galvo scanning (raster/spiral) to scan a diffraction limited *spot* over the cell body and; (2) **parallel** excitation which relies on light sculpting techniques (digital holography/temporal focusing) to generate a *blob* that simultaneously excites the entire cell body volume.

Importantly, both techniques can take advantage of combinations of fast galvo hopping to activate several neurons in quick succession, digital holography to generate duplicate spots/blobs targeting multiple neurons simultaneously, and temporal focusing to restrict the axial extent of the excitation volume. They also come with trade-offs dictating which experiment types they will be useful for. These relate to how they differ in terms of excitation volume size, laser power and time required for effective activation, as well as the mode of potential tissue damage that they risk.

Assuming that the serial excitation volume (a) is 0.25 µm^2^, the parallel excitation volume (A) is 100 µm^2^ and the ratio between the power required for parallel over serial excitation is √(A/a) (Peron and Svoboda, 2011), then serial activation requires an estimated 20 times less power and a 400 times smaller excitation volume than parallel excitation, at the cost of time spent scanning. In practice, differences in observed power requirements are likely to be affected by saturation of excitation in the focal volume, which is associated with efficient out-of-focus excitation as shown experimentally (J. Rickgauer and Tank, 2009). The degree to which differing photostimulation strategies (parallel versus serial) generate out-of-focus excitation has yet to be fully quantified and would be a useful direction for future research. Nonetheless, this lower requisite power means more neurons can be targeted simultaneously, however the necessity to scan results in potentially longer and more variable delays between excitation and action potential generation. While both strategies have the potential to cause tissue damage, the smaller serial excitation volume, and associated greater peak power, is more likely to generate damage via non-linear two-photon effects whereas the larger parallel excitation volume, and higher requisite average power, is more likely to generate heating damage. Given that heating is a concern with both photostimulation strategies, energy delivery to the tissue (and thus photostimulation time) is also a major factor. This is particularly important for serial strategies which require time spent scanning.

Therefore, serial excitation strategies are more suitable for experiments where many neurons need to be simultaneously activated per stimulus epoch, or on systems where power is limited. Conversely, parallel excitation strategies are more suited to experiments requiring exquisite temporal fidelity of action potential generation. In both conditions, stimulation power and frequency must be moderated to limit tissue damage.

**Box 1.**
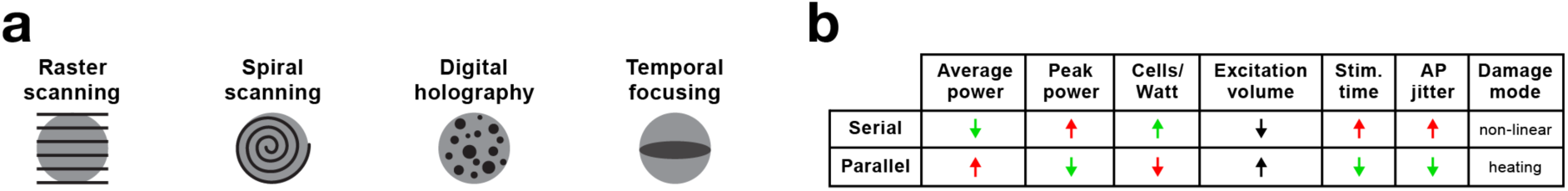
a. Schematic illustration of different photostimulation strategies b. Comparison of serial and parallel excitation methods (see Box 1 text for definitions). Favourable comparisons indicated in green, unfavourable in red.

##### Box 2 Two photon excitation and calibration of safe and effective stimulation parameters

Two-photon excitation is a nonlinear process whereby two photons must be absorbed nearly simultaneously by a molecule for it to be excited to a higher-energy state. This only occurs with a high probability when light intensity is high, such as at the spatial focal point of a beam focused through a lens. Lasers used for two-photon absorption are typically pulsed in time to enable light intensity to be concentrated in each individual pulse, increasing the probability that two-photon excitation events occur. Apart from wavelength, such lasers are typically characterized by three key parameters central to their ability to generate two-photon excitation: time averaged power, the width of the pulses in time (pulse width), and the frequency of the pulses (repetition rate). The probability of two-photon excitation is proportional to:

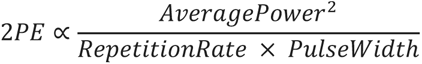

To photostimulate more cells more strongly one approach would be to increase the average power delivered to each cell. However, increasing average power will lead to thermal heating of the tissue and ultimately thermal damage (Podgorski and Ranganathan, 2016). Another approach could be to increase the intensity of light per pulse but keep the average power constant by reducing the pulse repetition rate. Note that reducing the pulse duration could also act to increase the intensity per pulse, but this will be associated with a concomitant increase in spectral bandwidth, which may cause chromatic aberration in SLM diffraction patterns as different wavelengths will be diffracted to different angles. We note that most if not all opsin (channel and pump) photocycles are longer (on the order of 10s of milliseconds (Zhang *et al*., 2011)) than the interval between laser pulses for even sub-MHz lasers. This is very different to fluorescence imaging, where the lifetime of the emitted fluorescence is much shorter than the interpulse interval. Opsins are thus ideally suited for use with for low-repetition rate, high peak power excitation (whereas fluorescence imaging necessitates high-repetition rates) as we and others have found (Yang *et al*., 2018). A typical microscope has a power throughput of ∼10% (dependent on number of elements and the optical coatings), meaning a typical 20 W laser will provide ∼2 W of power on the sample, potentially enough to stimulate ∼300 neurons at once. Typically we stimulate no more than 100 neurons at once for a short duration to mitigate heating concerns.

While using low rep-rate lasers reduces the risk of thermal damage (which is proportional to average power) increasing the probability of two-photon excitation does come with an increased risk of non-linear damage such as formation of intracellular plasma leading to ablation or lesioning of organelles (Koester *et al*., 1999; Hopt and Neher, 2001). It is therefore vital to carefully titrate the exact combination of average power, pulse width, and repetition rates to be used for experiments. The activation of neurons depends on the total deposited energy, but there may be different photodamage effects associated with how that energy is deposited. We typically test a series of increasing average powers at a given laser setting regime (wavelength, pulse width, repetition rate), and then look for successful stimulation responses as well as signs of damage such as increased baseline fluorescence. We then select the condition that provides successful stimulation without causing any sign of photodamage. The optimal combination of parameters will differ based on the above-mentioned laser parameters (average power, pulse width, and repetition rate) as well as wavelength, numerical aperture of the optical system, exposure duration and stimulus repetitions. The amount and type of non-endogenous protein expressed as well as the type of cell being tested is also likely to factor into whether the stimulated neurons respond with action potentials. Note photodamage, and photoablation are undesirable for most experiments, but some experimental paradigms use 2-photon ablation of specific neurons as a powerful manipulation (Peron *et al*., 2020).

**Box 2.**
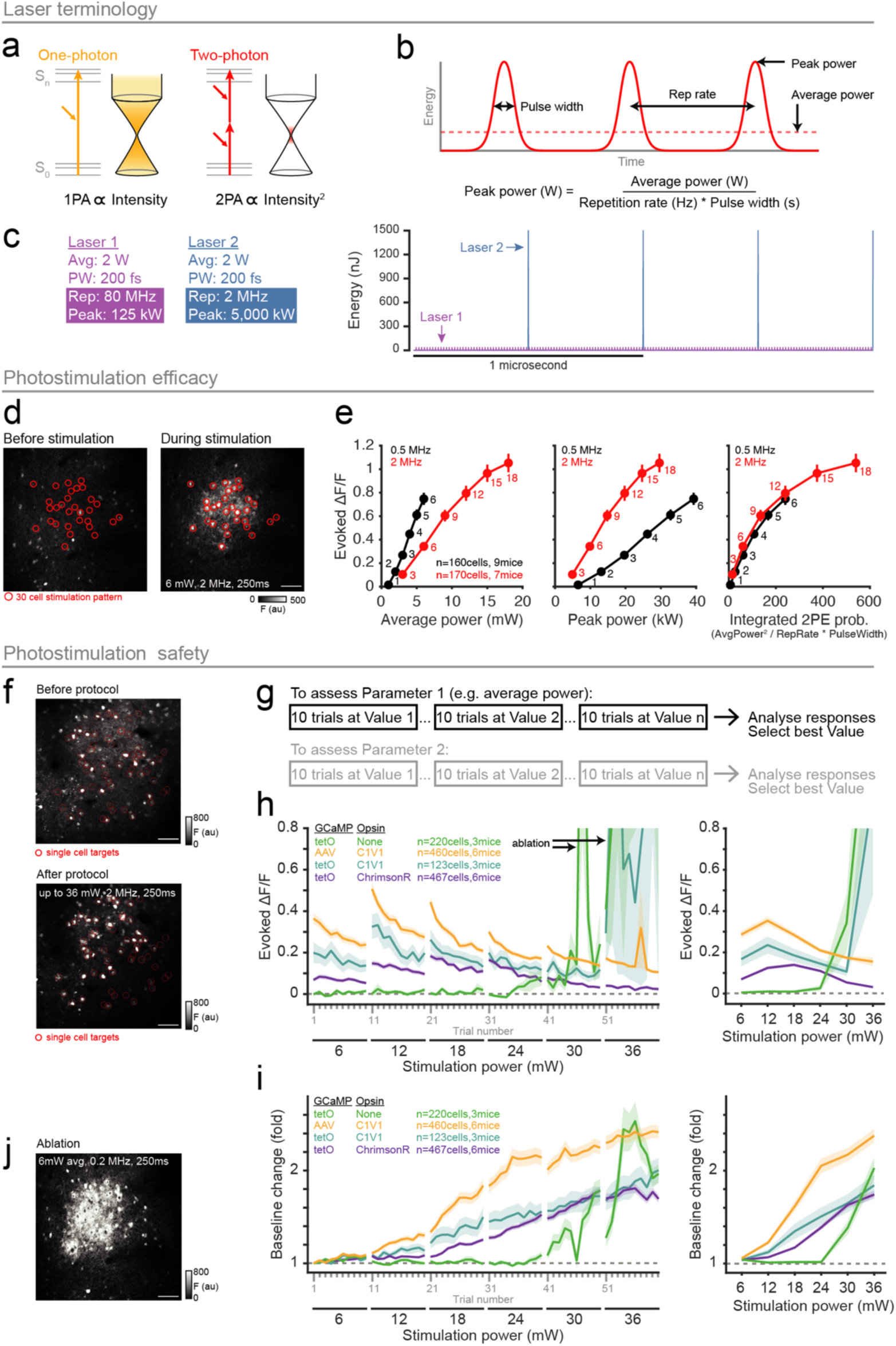
a. Two-photon excitation compared to one-photon excitation. b. Diagram illustrating key parameters of pulsed lasers used for both imaging and photostimulation applications. c. Hypothetical comparison between two lasers with the same average power and pulse width, but with a different repetition rate (and thus peak power). d. FOV fluorescence image before and after photostimulation of a 30-cell ensemble. Animals were wildtypes virally expressing AAV1-hSyn-GCaMP6s and AAVdj-CaMKII-C1V1. Scale bar represents 100 μm. e. Stimulation of 10-cell groups (pre-filter for responsive neurons) with range of average powers at two different laser repetition rates. These group were stimulated in blocks of increasing powers at one of the two repetition rates. Spiral parameters were as follows: 15 μm diameter, 20 ms duration repeated 5 times at 20 Hz. 10 trials of each stimulation were conducted, with 10 seconds between each stimulation. Plots show the average response size (ΔF/F) of the targeted ROIs as a function of either the average power per cell (measured on sample), the peak power, or the integrated probability of two-photon excitation. Numbers indicate the average power (mW). Based on these assessments 6 mW per cell (at 2 MHz) was chosen as an effective power to use for our stimulation experiments. f. To assess phototoxicity we stimulated single cells with a range of increasing average power (at 2 MHz rep rate). Cells were selected from GCaMP expression images with no knowledge of opsin expression. Various different expression strategies were used. tetO refers to transgenic animals expressing GCaMP6s under the tetO system (Wekselblatt et al. 2016). Spiral parameters were as follows: 15 μm diameter, 20 ms duration repeated 5 times at 20 Hz. 10 trials of each stimulation were conducted, with 10 seconds between each stimulation. Images show FOV of GCaMP expression before and after the single-cell stimulation protocol. Scale bar represents 100 μm. g. Outline of the calibration protocol. We select one parameter at a time keeping all others constant. We perform 10 trials at a given value of that parameter, and then increment it and acquire 10 more trials, repeating until we reach the maximum value to be tested. Subsequent analysis is used to select the power that was effective (resulted in activation of neurons) and also safe (no changes in baseline fluorescence). If there are multiple parameters to be tested we would then proceed in a similar fashion with the next parameter. h. Average response size of the targeted cells across trials and blocks of increasing power. Responses all tend to decrease within a block, likely due to opsin desensitization. The large ‘responses’ at 30 and 36 mW are a result of photoablation. i. Average baseline fluorescence of the targeted cells across trials and blocks of increasing stimulation power. We used the baseline fluorescence as an indicator of cell health, with increases (that were not the tail end of GCaMP transients) representing an undesirable change. Note the strong increase after a few trials at 18 mW. j. An example of intentional photoablation of a multiple cell ensemble, using a very low (0.2 MHz) repetition rate laser.

### Expertise needed to implement the protocol

The setup, maintenance and use of all-optical microscopes requires considerable optical expertise. Primary scope users should have the responsibility of maintaining and calibrating the microscope on a daily basis. They should be familiar with the full light-path alignment procedure and they should be experienced in diagnosing and correcting misalignments. Beyond this, any user of the protocol described below requires minimal expertise beyond the animal handling required for the particular experiment being carried out, and an ability to operate basic imaging and photostimulation functionality in the microscope software.

### Limitations

We have successfully applied this protocol for all-optical experiments using commercial Bruker (Russell *et al*., 2019; Dalgleish *et al*., 2020) and Thorlabs (Robinson *et al*., 2020) microscopes. While the general principles will apply to any setup configured for all-optical experiments, our software routines (available with this protocol) may have hardware-specific implementations which may preclude their use with other systems.

## MATERIALS

### REAGENTS

- Mice prepared for chronic all-optical experiments (expressing both an indicator and an opsin with non-overlapping excitation spectra, implanted with a chronic window over the expression site) **! CAUTION** All animal experiments must comply with the relevant institutional and national animal care guidelines.

○ Wildtype or transgenic (expressing indicator, opsin or a recombinase) mice
○ Viruses encoding the indicator and/or opsin, as well as dilution buffers to achieve suitable concentration for injection
○ Headplate
○ Chronic optical window and/or a cannula (for deep preparation)
- If anaesthesia is required: anaesthetic (e.g. Isoflurane) and eye lubricant (e.g. Allergen Lacri-Lube, Allergen Pharmaceuticals, Ireland)
- Fluorescent plastic slide for calibration routines (Chroma, Thorlabs)
- Immersion fluid if using immersion objectives (e.g. distilled H_2_O)
- Sucrose water for behavioural experiments

### EQUIPMENT

- Surgery setup requires a pipette puller and beveller, injection syringe pump with stereotaxic control and microscope. The animal is anesthetized and held in place by ear bars prior to implantation of a headplate. For preparations involving deep brain areas a cortical aspiration setup may be necessary.
- Two-photon all-optical microscope for *in vivo* imaging and SLM-based photostimulation (Bruker, Thorlabs, Scientifica, 3i, custom build, etc.)
- Objective suitable for two-photon imaging (e.g. Nikon 16x/0.8-NA, Leica 25x/0.95-NA, Thorlabs 10x/0.5-NA)
- Imaging laser suitable for two-photon excitation of the indicator and opsin-fluorophores (e.g. a tunable Ti:Sapphire high repetition rate laser such as a Coherent Chameleon or a SpectraPhysics MaiTai capable of 920 nm and 765 nm for our combination of GCaMP6s and C1V1-Kv2.1-mRuby respectively).
- Photostimulation laser suitable for two-photon activation of opsins (e.g. Amplitude Satsuma, Coherent Monaco, Menlo BlueCut, SpectraPhysics Spirit) **! CAUTION** two-photon lasers used for photostimulation tend to have very high average powers (>5 W) and pulse energies (>10 µJ) and can cause serious burns and eye damage as well as damaging equipment. Always comply with laser safety regulations and ensure that all elements in the photostimulation light path have the requisite optical power tolerance.
- Microscope control software (PrairieView for Bruker systems, ThorImage for Thorlabs systems, ScanImage for Scientifica systems)

○ MATLAB (>2016a) for custom calibration, setup and analysis programs
○ Naparm control software (https://github.com/llerussell/Naparm)
○ STAMovieMaker software (https://github.com/llerussell/STAMovieMaker)
○ TransformMaker (https://github.com/llerussell/SLMTransformMaker3D)
○ PhaseMaskMaker (https://github.com/llerussell/SLMPhaseMaskMaker3D)
○ RawDataStream (https://github.com/llerussell/Bruker_PrairieLink
- Head fixation apparatus and animal holding platform/treadmill and a soundproof enclosure for controlled behavioural experiments
- Behavioural control software and hardware (e.g. PyBehaviour https://github.com/llerussell/PyBehaviour) with lick and motion detectors, stimuli presenters and reward delivery
- Software for embedding two-photon and one-photon optogenetic stimulation patterns into behavioural paradigms (e.g. Two-Photon Behaviour Sequencer https://github.com/hwpdalgleish/TPBS)
- DAQ cards (e.g. National Instruments) and synchronization software (PackIO for most systems www.packio.org, or ThorSync for Thorlabs systems)

## PROCEDURE

### Module 1 (Figure 4) – Calibration of all-optical system

A critical step in successful all-optical experiments is achieving accurate targeting of the two-photon photostimulation laser to neurons identified with two-photon imaging **(Figure 4a)**. The first step is ensuring the two laser paths are physically co-aligned as much as possible by adjusting mirrors to hit established alignment targets. Minor corrections to the co-registration can be made by applying an offset to the galvanometer mirrors specific to the photostimulation pathway, ensuring that both beams point to the centre of the FOV. However as the two optical pathways are independent they still operate in different coordinate systems. When using an SLM, the mapping from SLM-space coordinates into imaging coordinates needs to be computed. To calculate this required transformation, prior to the actual experiment we focus the photostimulation laser into arbitrary spot patterns to burn holes in a fluorescent plastic slide (Chroma, Thorlabs), then by imaging the same area with the imaging laser we can register the intended (programmed) locations of the burns with the actual achieved location of the burns in imaging space **(Figure 4b-d)**. Note that the burnt spots are not necessarily parfocal with the imaging plane and therefore a stack of the plastic slide is acquired. This registration is well captured by an affine transformation (a geometric transformation that preserves collinearity) between the two coordinate systems and results in a mapping from SLM coordinates to imaging coordinates, enabling us to program the SLM to target identified neurons in the imaging FOV **(Figure 4e-f)**. Co-alignment (steps 12-14 below) should be checked often to ensure accurate targeting in all experiments. We recommend checking this often (e.g. daily) in the first instance to confirm there are no drifts in the system but eventually less often - e.g. weekly to monthly if there is sufficient temperature stability in the room and pointing stability of the laser source, both of which are required for consistent optical alignment. The procedure to align and calibrate the two lightpaths is as follows:

1. Centre galvos in both imaging and photostimulation pathways.
2. Ensure adequate physical alignment of the photostimulation beam through the entire pathway up until the SLM, hitting alignment targets at various manufacturer design points. The beam should overfill the active surface of the SLM to maximise optical resolution, especially where power throughput is not a major concern.
3. Unblock the zero order of the SLM by translating the block (see above) out of the way, and continue to optically align through the system with this beam again hitting the manufacturer alignment points, culminating at the back aperture of the objective. Both beams should hit the centre of the back aperture. Depending on the magnification factor the photostimulation beam may be smaller than the imaging beam, and the appearance of the rectangular reflective surface of the SLM may be visible.
4. The SLM’s efficiency is in part dictated by the polarization of the laser beam. The polarization of the beam can be adjusted with a half wave plate on the optical table. One way to do this is as follows. Position a fluorescent card at the zero order block position to visualise the diffraction pattern. Apply an arbitrary diffraction pattern and optimize the power distribution into the first order spots, and out of the zero order spot, by rotating the half wave plate. After optimization, remove the fluorescent card.

- Other methods to more accurately optimize the polarization could be to either focus the SLM spot pattern on a fluorescent slide under the objective and visualise the pattern with a camera and repeat the procedure using pixel intensity to quantify the ratio of zero order to first order brightness. Another method would be to position a power meter under the objective - after having blocked the zero order - and optimize the polarization to reach maximum power intensity, which corresponds to the optimum first order diffraction.
5. Now that the photostimulation beam is physically coaligned with the imaging pathway, position a plastic slide under the objective.
6. Ensure parfocality of the imaging and photostimulation beams. Upload a phase mask to the SLM and burn the spot pattern in the plastic slide. Find the axial centre of the burn location by moving the imaging z-focus. One way to fine-tune the parfocality is to make slight adjustments to the vergence of the stimulation beam (assuming the imaging pathway is well collimated) by adjusting the second lens of the post-SLM telescope to fine-tune the parfocality until the burn location is at the nominal imaging plane.

- Burn parameters: 50 mW per spot, 10 ms duration, repeated until burns are visible (Amplitude Satsuma 20 W 2 MHz).
7. Remove the phase mask on the SLM by uploading a blank phase mask, so that only the zero order beam is propagated through the system.
8. Decide on the optical zoom of the imaging pathway to be used for experiments as the photostimulation calibration is specific to particular imaging conditions.
9. Perform the manufacturer’s procedure to align the galvanometer pointing of the photostimulation pathway (using the unblocked zero order beam) with the imaging pathway.

- Burn parameters: 50 mW, 10 ms duration, repeated until burns are visible (Amplitude Satsuma 20 W, 2 MHz).
10. Now, re-block the zero order by translating the block into place. This is straightforward to achieve when using a camera to image the spot patterns on a slide.
11. Map out the zero-order blocked region. Apply a grid-like spot pattern and visualise the fluorescence with a camera in order to calculate the size of the block in physical space. This region of SLM space is unaddressable, but note that by translating the galvo pointing position the blocked region can be avoided.
12. Next, the SLM targeting calibration is performed. Display an arbitrary, rotationally asymmetric pattern (with no transform applied) on the SLM and burn spots in the plastic slide. If performing a 3D calibration burn this pattern at various axial offsets using 3D phase masks. The arbitrary SLM Z-coordinate range can be found by trial-and-error until the burns are within the desired volumetric imaging range. Take an image (or a stack if performing a 3D calibration)

- Burn parameters: power 50 mW per spot, 10 ms duration, repeated until burns are visible (Amplitude Satsuma 20 W, 2 MHz).
13. Inspect the acquired image and record the coordinates of the burn locations in the image and use these to compute the affine transformation between the intended (i.e. programmed SLM coordinates) and the actual burn locations. See SLMTransformMaker.m for an example implementation.
14. Generate new spot patterns – with the newly calibrated transform applied – and again burn spots in the slide. Take an image (or 3D stack) and calculate the distance from intended targets to the actual burns to test the accuracy of the calibration. Redo steps 12 & 13 if necessary, e.g. if the burnt spots are greater than 2 µm from the intended targets (acceptable accuracy will depend on the structures being targeted)
15. Finally, the total power throughput (i.e. the average power on sample) is calibrated to ensure safe and effective stimulation of neurons. Apply a typical SLM spot pattern (i.e. one similar to what might be used during an experiment) and measure power on sample after the objective with galvos centered. Record the power at intervals of the power modulation device setting. This number is used later in combination with the number of intended neuron targets to set the total power level (split between all the targets).

- Optional: To calibrate the reduction in efficiency with larger diffraction angles displace a single point to increasing offsets from the zero order and record the power at each location. The relationship between distance and power can be used to scale the power distribution amongst spots in order to equalise the power delivered among spatially dispersed neurons in a group.

### Module 2 (Figure 5) – Surgery TIMING 3 hours

The target neurons must be engineered to express two proteins which are typically delivered virally (Packer *et al*., 2015). To provide optical access to expressing brain tissue the overlying skull is replaced by a chronic window (Holtmaat *et al*., 2009), and for deep structures some superficial brain tissue may need to be removed (Dombeck *et al*., 2010). A headplate is installed to enable head-fixation under microscopes.

**! CAUTION** Ensure relevant regulations and guidelines for sterile recovery surgeries are followed at all times.

1. Determine the best expression strategy for desired experimental protocol (**Figure 5a**), taking into consideration viral injection volumes and concentrations if using virally expressed constructs, and the complexity of combining genotypes if using transgenics.
2. By definition all experiments will require optical access to the brain region of interest. Ensure that the optical setup of the microscope, surgical preparation and spectral properties of all constructs are appropriate, particularly when the brain region of interest is deep (**Figure 5c,d**). Possible optimizations could be to use far red-shifted indicators and/or opsins to reduce scattering of excitation photons, or methods to improve access to deep structures (i.e. GRIN lenses, cortical aspiration).

▴ **CRITICAL STEP** AAV serotypes and promoters should be chosen to ensure specificity of expression in the cell population of interest while also allowing sufficient overlap of opsin and indicator expression (see **Module 3a, Step 5** for more detail).

▴ **CRITICAL STEP** Ensure that there is sufficient spectral separation between the 2- photon excitation spectra of opsin and indicator to avoid cross-talk in either direction. The imaging laser wavelength should not significantly increase spiking in opsin-expressing neurons (Packer *et al*., 2015), nor should the photostimulation laser wavelength induce appreciable fluorescence of the indicator (although note that photostimulation lasers may also cause significant, and unavoidable, autofluorescence of endogenous fluorophores in the tissue). In practice, photostimulation can cause imaging artefacts. However, given that these are limited to the exact time of photostimulation, and most calcium indicators have a slow decay, this may not be an issue in practice if photostimulation epochs can be discarded and responses can be analyzed in the post-stimulus window.

▴ **CRITICAL STEP** Ensure that the expression strategy yields robust expression in sufficient numbers of neurons for your experimental purposes without damaging the region of interest. If using viral strategies, make sure that the total volume of virus injected and the number and proximity of injection penetrations to cortical site of interest does not damage surrounding tissue. If using transgenic strategies, which tend to drive expression in fewer neurons than virally-mediated expression, make sure that any transgenic lines used yield sufficiently dense expression (see **Module 3a, Step 5** for more detail).

3. Decide on surgery protocol, using appropriate co-ordinates, to target your brain area of interest (**Figure 5b – d**) and perform the surgeries. **? TROUBLESHOOTING** **! CAUTION** Allow sufficient recovery time between surgery and subsequent procedures according to the guidelines of your institution.
4. If using viruses, allow sufficient time for constructs to express to useable levels (∼2 – 3 weeks for AAVs). If using transgenics, ensure that surgery is done at an appropriate time in the mouse’s life-cycle such that the transgene of interest is robustly expressed. See **Module 3a** below for details on identifying useable expression.

### Module 3a (Figure 5) – Visualizing opsin and indicator expression TIMING 10-20 min

1. After having allowed animals to express constructs for a sufficient time (see above), headfix the animal beneath the microscope. If anaesthesia is required then first anaesthetise in 5% isoflurane in an induction chamber, then transfer to the two-photon microscope, maintaining anaesthesia with 1% isoflurane and mouse body temperature with a heating blanket. **! CAUTION** All animal experiments must comply with the relevant institutional and national animal care guidelines.
2. Navigate to the relevant FOV.
3. Take anatomical image(s) of plane(s) of interest to check expression of opsin (e.g. at 765 nm for C1V1-Kv2.1-mRuby) and indicator (e.g. at 920 nm for GCaMP6s) (**Figure 5b – d**).
4. Ensure that expression is sufficient for your experimental purposes. Typically this should be as robust as possible (not under-expressing) without causing neuronal damage (not over-expressing). Specifically, cytosol-filling indicators (such as GCaMP) should show low baseline fluorescence and high SNR transients with few neurons having filled nuclei (<10% of neurons; Tian *et al*., 2009; Chen *et al*., 2013; Packer *et al*., 2015). Opsin-conjugated fluorophores (if present) should be clearly visible in appropriate cell compartments (i.e. only in the soma if using soma-restricted opsins) with reasonable laser power (<50 mW) (**Figure 5b – d**), and any 2-photon photostimulation should yield reliable calcium transients (see **Module 4b, Step 13** below for details) in targeted neurons without excessive recruitment of off-target neurons (i.e. significant activation of neurons outside of the measured resolution of the system that decays with distance from stimulation sites). TROUBLESHOOTING

▴ **CRITICAL STEP** Ensure that both opsin and indicator express in enough neurons in the neural population of interest and that there is sufficient overlap between them. The requisite number of construct-expressing neurons will vary, but for our experiments using all-optical techniques to modulate behavior we typically use FOVs in which we have identified >150 opsin-expressing neurons and >500 indicator expressing neurons in a given 710 µm imaging plane of our 4-plane volumetric stack (33 µm spacing; 100 µm axial extent) (Dalgleish *et al*., 2020), corresponding to a pool of >600 opsin and >2000 indicator expressing neurons across the volume. Ideally overlap between opsin and indicator would be 100%, as it is when using bicistronic opsin-indicator constructs (Marshel *et al*., 2019). However, in our experiments using separate viral opsin and indicator constructs, or viral opsin in indicator transgenic mice, we routinely use FOVs where only 40 – 50% of indicator-expressing ROIs also have opsin, corresponding to >50 dual expressing neurons in a given FOV as described above (Dalgleish *et al*., 2020) and therefore >200 total dual expressing neurons across the volume to choose from for all-optical interrogation.

▴ **CRITICAL STEP** Always use similar laser power for expression checking (30 – 50 mW for cortical depths of 100 – 300 µm) to ensure that differences in apparent expression clarity/brightness are due to the expression itself and not variation in strength of fluorophore excitation.

### Module 3b (Figure 6) - Training animals on a behavioural task TIMING 7-10 days

To probe the neural basis of a perceptual or behavioural function we require the animals to perform a reliable and repeatable behaviour. Typically these are tasks whereby the animal indicates the presence of a particular stimulus, in most cases by licking at a water spout in order to receive a sugar water reward for the correct response. Mice are motivated to perform the tasks by being placed on a food or water restriction diet and learn the task through a series of iterative steps (Guo *et al*., 2014). The time taken for animals to reach good performance on the task will depend on the complexity of the task.

1. Train animals on desired behavioural task, following the normal habituation, learning and training phases (**Figure 6**; see below).

▴ **CRITICAL STEP** Choose an appropriate behavioural task design (Carandini and Churchland, 2013) for your experimental question as this strongly dictates the types of all-optical manipulations possible and the causal inferences that can be drawn. For instance, in our lab we routinely use: detection tasks to assess how the number and functional identity of stimulated V1 neurons influences the detection threshold of mice trained to detect orientated gratins (**Figure 6a**); discrimination tasks to assess how much additional activity in S1 is required to bias animals to choose stimulation of one whisker over another (**Figure 6b**) and how this is integrated into local network processing of sensory information; complex behavioural tasks, such as head-fixed navigation along virtual linear tracks, to compare how targeted place cell perturbations influence local network activity in hippocampal CA1 and CA3 (**Figure 6c**).

▴ **CRITICAL STEP** Ensure that the task can be learned in a timeframe such that the time when animals perform at desired levels coincides with the period during which constructs are optimally expressed (i.e. if using virally-mediated constructs before constructs begin to overexpress and degrade cell health; if using transgenic animals, then after transgene expression has begun and before it ceases). If this presents a problem, consider a two-stage surgical protocol where initial installation of a head-fixation device allows prior training to the desired level of performance before subsequent expression of constructs and chronic window installation.

### Module 4a (Figure 7) – Mapping functional properties of neurons TIMING 1-2 hrs

1. Headfix the animal beneath two-photon microscope. If anaesthesia is required then first anaesthetise in 5% isoflurane in an induction chamber. Then transfer to the two-photon microscope, maintaining anaesthesia with 1% isoflurane and mouse body temperature with a heating blanket. **! CAUTION** All animal experiments must comply with the relevant institutional and national animal care guidelines.
2. Find and map the FOV in your brain region of interest (**Figure 7**). An appropriate protocol for this will be similar to most functional mapping experiments used in correlational studies, however it should be optimized to allow fast online analysis (**Figure 7a**). We will briefly describe an example below.

▴ **CRITICAL STEP** Online analysis allows functional mapping to inform subsequent all-optical interrogation even within the same experimental session without the animal becoming too unmotivated and/or tired to perform. The major optimization we have found useful is to register our 2-photon time-series in real-time by streaming raw pixel data from microscope acquisition software directly through a custom pipeline in MATLAB (RawDataStream; https://github.com/llerussell/Bruker_PrairieLink). This reduces post-acquisition time to register the data from ∼1 minute/minute of acquired data per plane to essentially zero. By writing the data to a raw binary file, readable by any programming language, we avoid the TIFF file format and the overheads associated with slow loading. After the data is acquired we use this instantly-registered data to identify targets in one of two ways. First, we generate pixel stimulus-triggered average (STA) images (using STAMoviemaker; https://github.com/llerussell/STAMovieMaker) in which the post-stimulus response of each pixel, averaged across trials, dictates the hue (preferred stimulus identity), saturation (preferred stimulus tuning strength) and value (preferred stimulus response amplitude) in the generated image. This gives a quick, intuitive map of the strength and tuning of functional responses that is automatically in register with the spatial position of the imaged neurons, providing the information needed to place photostimulation targets at the spatial location of functionally tuned neurons of interest (see below and **Fig 7.b,c,e,f,h,j** for examples). Alternatively, when more sophisticated analysis of responses is required, we use a version of the Suite2p toolbox (Pachitariu *et al*., 2016) modified to read the raw real-time registered time-series binary files, skipping the lengthy registration step to generate ROIs and traces in ∼5 mins/plane (+ 5 mins for manual ROI curation). This allows us to confirm the intuitive results of STA image analysis and generate target groups on the basis of statistical comparisons between neuronal traces.

3. Set up stimuli to map, e.g. visual gratings for V1 (**Figure 7b – d**), whisker vibrations for S1 (**Figure 7e – g**), or navigation in virtual environments for hippocampal CA1 and CA3 (**Figure 7h – j**).
4. To find a 2-photon FOV for all-optical interrogation that has the functional responses desired and robust, healthy construct expression (**Figure 7b,e**), use STA images from functional widefield calcium imaging and structural construct expression images (see **Module 3a**).
5. Map the functional responses of the desired FOV at the cellular level using 2-photon imaging, generate 2-photon STA images (**Figure 7c,f,h,i**) and use online ROI/trace analysis to extract trial-wise traces (**Figure 7d,g,j**; left) which can be used to extract average functional tunings (**Figure 7d,g,j**; right). TROUBLESHOOTING

### Module 4b (Figure 8) – Mapping photostimulation response of targeted neurons TIMING 30 min

Before performing all-optical experiments it is beneficial to know which neurons are photostimulatable – i.e. identify those neurons which express both the indicator and opsin to sufficient levels to enable optogenetic activation while their activity is recorded. Baseline marker fluorescence is insufficient to ensure functional expression levels. Therefore, to identify these cells we photostimulate each and every cell in the FOV and record their responses. This could be achieved by stimulating single cells, one by one, but this becomes time-consuming when the number of cells in the FOV is high. A second option would be to photostimulate all cells at once, but this is only appropriate if the number of cells in the FOV is low (due to a limited laser power budget, as well as concerns over heating). An optimal solution is to group all the neurons into a number of groups of a defined size and stimulate the groups one-by-one. We have designed a piece of software (Naparm, Near-Automatic Photoactivation Response Mapper) that implements this stimulate-all-cells ‘mapping’ protocol as well as being flexible enough to implement any other type of all-optical experiment where only a certain group(s) of selected neurons are stimulated. This software makes it intuitive to identify ROIs in a FOV, and design stimulation group and parameters and then generate all the files required to configure the microscope to execute the experiment.

1. Run Naparm and import relevant FOV image(s) by either dragging images into the image window (left) or clicking the *Load image(s)* button in the **Image** panel. For standard photostimulation response mapping this will likely be the opsin image(s). Note that multiple image(s) of the same plane(s) can be imported in this way and viewed via the drop-down menu in the **Image** panel.
2. Select opsin-expressing neurons to photostimulate. Either do this manually by left-clicking on cells in the image window (left), or use one of the automatic detection options in the *Automatic* section of the **Add points** panel.

- Optional. Note that by applying a grid of equally spaced points over the whole FOV in lieu of precisely targeted neurons, a good estimate of neuronal photostimulation responses can still be obtained without the time-consuming step of identifying hundreds (or thousands) of neurons (see **Figure 8i**).
3. Once all relevant neurons have been selected, group them into stimulation groups (cells within a group will be stimulated simultaneously, and groups will be stimulated sequentially). Select the desired grouping method from the **Assign groups** panel and adjust the *Group size* or *Number of groups* options to desired value. For standard photostimulation response mapping we use the *ekmeans* algorithm to group neurons into spatially clustered groups of equal size. Click *Group* to group cells and note that selected cells in the image window are now colored by group and associated with the centroid of all cells within their group. For randomly seeded algorithms (*ekmeans* and *Random*) clicking *Group* multiple times will repeat the grouping procedure and reassign cells to groups. In general we use a group size of 50, grouped via the *ekmeans* algorithm.
4. Set up the timing structure of how and when groups are stimulated within a single trial of the protocol; this will define the stimulation pattern both for a given group, and how the groups are sequenced through. In the **Single trial** panel, for a single group set the number of times to stimulate it, at what rate and with what photostimulus duration with the *Shots per pattern*, *Inter shot interval (ms) (i.e. the timing between spiral onsets)* and *Spiral duration (ms)* fields respectively. Use the *Delay first spiral (ms)* to define the time it takes for your SLM to update to a new pattern. For our system using a BNS P512-1064 SLM we use 5 ms (For our system using a Meadowlark P1920 SLM we use 20 ms due to slower pixel response times). Define the time between each group with the *Change pattern every (ms)* field. The *Trigger each pattern* checkbox to decide whether to trigger each pattern individually (checked) or just the first pattern (unchecked). Note the latter option assumes that the sequence of patterns will be generated by some external software, e.g. the microscope software itself (as in the Bruker system). The entire sequence of groups can then be repeated on a given trial by setting the *Sequence repetitions* and *Sequence repetition interval (ms)* fields. Note that all of these fields will dynamically update the trigger display below. Colors in this display correspond to group colors in the image window (left). In general we stimulate 50 cell patterns (total number of patterns depends on the total number of targets in the FOV) with 10 shots at 20 Hz (50 ms inter-shot intervals) with a spiral duration of 20 ms and a 5 ms delayed first spiral. We sequence from one pattern to the next every 1 second.
5. Set up the timing of how many and how often single trials are delivered in the complete protocol. In the **All trials** panel update *Number of trials* to define the number of repeats of the single trial defined above. Set the inter-trial interval with the *Trial length (s)* field. Note that this is inclusive of the time it takes for the sequence of stimuli to be stimulated. If appropriate, add a period of spontaneous imaging (i.e. no photostimulations) either before and/or after the photostimulation mapping period. Again note that changing these fields will dynamically update the trial triggers display below. In general we do 10 trials, separated by a minimum of 10 s inter-trial interval.
6. Set the parameters of how the neurons are stimulated by the laser, in this case in terms of power and spiral shape/size. In the **Spiral parameters** panel, the *Revolutions* field defines the number of revolutions that describe the spiral shape itself. To repeat a given spiral multiple times, change the *Shots per pattern* field in the **Single trial** panel. The *Laser power* field defines the power distributed across all targets in a given group. In general we use 3 spiral revolutions, a 15 µm spiral and a laser power that provides 6 mW per cell. **! CAUTION** Ensure that your laser power is calibrated, so that the value entered into Naparm corresponds to the power on sample. Also make sure to test safe laser powers per cell for your desired photostimulation pattern and be careful not to exceed this (see **Box 2**).
7. Depending on the FOV size (imaging and stimulation), dispersion of the neurons within and between the groups and the diffraction efficiency of the SLM decide whether to use galvo/SLM (whereby the galvos are steered to the centroid of each group and the SLM patterns are relative to this new set point) or pure SLM targeting with the *Centroids* and *All points* buttons respectively in the **Mark Points mode** panel.
8. Now that the experiment is designed, export all necessary files in order to then load them into and configure the microscope software. In Naparm export these files using the *Export all* button at the bottom right of the GUI. This will output a folder with a user-defined name into a user-defined path (path and name are defined earlier by the “*…”* button and text box in the **Save path** panel) containing the requisite files for the microscope system. These files should be loaded into the relevant sections of the microscope control software.

○ This folder will contain a file with the photostimulation galvo locations (.gpl file for Bruker systems, .bmp for Thorlabs systems), an .xml file, and a folder of phasemasks. The .xml defines the photostimulation protocol (timing, spiral parameters and photostimulation power).
○ If using external software to upload the phasemasks to the SLM (i.e. not driven by the microscope software) then the folder of phasemasks will contain all phasemasks used to complete a single repetition within a single trial. Load this folder of phasemasks into your SLM control software (in our case Blink with OverDrive Plus) and set the number of repetitions to a value appropriate for your protocol.
○ The .dat files contain triggers for the SLM updates (changing patterns) and spiral delivery. These should be loaded into the master synchronization software (in our case this is PackIO).
9. Setup an imaging time series (t-series) acquisition of the plane(s) of interest with the requisite number of frames. Allow a buffer of ∼10 s before and after the photostimulation mapping protocol for pre- and post-photostimulation analysis.
10. Check via an imaging “live scan” (i.e. not the full acquisition) that the imaging FOV has not moved significantly in the time it took to setup the photostimulation mapping protocol. To do this find an obvious landmark (e.g. a cell highly expressing indicator) in your anatomical image(s), note the pixel location of that cell’s centroid and ensure that the cell is at that location in the live imaging window. If using a water immersion objective, also confirm that the immersion fluid level is sufficient. Stop the “live scan”. **! CAUTION** If the FOV has moved by > 5 µm then photostimulation targets will no longer efficiently target the desired cells.
11. Begin the photostimulation experiment. Arm all relevant software to wait for triggers. Begin the master synchronisation software recording. Begin the t-series. Wait 10 s, then begin delivering the photostimulation triggers from the synchronisation software.
12. Once imaging is complete, analyse the results of the photostimulation experiment. One method to do this is via the construction of pixelwise STA images, revealing which neurons were successfully stimulated (**Figure 8g, i**) (note that another method is to use extracted traces and this is described in **Step 13** below).

- Import both the t-series file and synchronisation file into STAMovieMaker. This will process and save out STA (stimulus-triggered average) movies and images for further analysis and cell selection.
- Configure the GUI for photostimulation STA analysis. Update the *Different stims* field in the **Stim setup** panel to the number of groups stimulated.
- Click Run and then inspect the output images and movies for brightly colored (successfully stimulated) neurons. TROUBLESHOOTING
13. When analysing extracted traces from imaging data, neurons responsive to photostimulation should show reliable calcium transients (transients on >50% of photostimulation trials) of an amplitude appropriate for your chosen opsin/indicator combination, photostimulation parameters and experimental requirements. For example, for our most commonly used indicator (GCaMP6s) we use the fluorescence change elicited by known numbers of spikes (Chen *et al*., 2013), confirmed with our own simultaneous calcium imaging and cell-attached electrophysiological recordings (Packer *et al*., 2015), to estimate the transient amplitude resulting from a single spike on our system (∼0.1 ΔF/F). We then confirmed this for photostimulation with our most commonly used opsin (C1V1) for a given spiral power and duration (Packer *et al*., 2015). We use this quantal value, in combination with C1V1’s ability to follow photostimulus trains at different frequencies (Prakash *et al*., 2012), to estimate the expected transient size resulting from any photostimulus train we might use. For instance, in most Naparm protocols stimulating neurons 10 times at 20 Hz, given a single spike amplitude of ∼0.1 ΔF/F and a 90% response reliability of C1V1 at 20 Hz, we would expect a transient of ∼0.9 ΔF/F. In practice, for most of our experiments we tend to accept neurons that reliably respond less strongly than this expected value (>0.3 ΔF/F on >50% of trials) because the specific number of action potentials elicited in each neuron is less important to us than the number of neurons that we can activate to any extent. The amplitude threshold will therefore be set by the requirements of each experiment. Given that there is known cell-to-cell variation in sensitivity to photostimulation (Mardinly *et al*., 2018; Dalgleish *et al*., 2020; Sridharan *et al*., 2021), it is possible to tailor the photostimulation power delivered to individual neurons in a targeted population on the basis of their photostimulatability to equalise evoked responses across all neurons stimulated by using an SLM to modulate the intensity of individual diffracted beamlets (Mardinly *et al*., 2018).

### Module 5 (Figure 9) – All-optical interrogation during behaviour TIMING 3-5 hours

As described above, the workflow of the experimental phase of most all-optical behavioural experiments will follow a similar general structure (**Figure 2, Figure 9a**). We detail a specific example of this below (**Figure 9b – h**). For this experiment, a mouse has been trained over the course of ∼5 days to report 1P photostimulation of neurons in barrel cortex of decreasing power by licking for sucrose at an electronic lickometer. At the lowest LED power it has then been transitioned to detecting 2- photon photostimulation of arbitrary groups of 200 and 100 neurons. In the final phase of the experiment described below we want to characterise the functional tuning and photostimulatability of neurons in the C2 barrel (**Figure 9b**) and identify an ensemble that responds strongly to both whisker stimulation and photostimulation (**Figure 9c – d**), embed this ensemble into our behavioural training paradigm as one would any other stimulus type (**Figure 9e**) and photostimulate them during the final sessions of this previously learned behavioural task to assess their behavioural salience (**Figure 9f – h**).

1. At least 1 day prior to the final training sessions, anaesthetise animal and headfix on heatpad beneath microscope (5% for induction, 1% for maintenance).
2. Map the C2 barrel (see **Module 4a**, **Figure 7**) by delivering 10 trials of 30 Hz sinusoidal vibration in the dorso-ventral axis, 1 s stimulus duration, with a 10 s inter-stimulus interval using a piezo electric bender attached to the whisker via a glass capillary while wide-field imaging the dorsal surface of the brain through the cranial window using a blue LED, 5x/0.1-NA air objective and a sCMOS camera acquiring at 10 Hz (∼1.5 mm x 1.5 mm, 512 x 512 pixel resolution).
3. Use the sCMOS/LED light-path on the microscope to centre objective over region of indicator/opsin expression visible through cranial window.
4. Feed the C2 whisker into a pulled glass capillary attached to a piezo bender and turn isoflurane to 0.5%.
5. Acquire a widefield movie while delivering whisker stimuli.
6. To find the best sensory-responsive FOV for subsequent experiments, use STA Movie Maker to create a stimulus-triggered average ΔF/F image of the post-stimulus epoch from wide-field movie and stimulus triggers. The peak in this image corresponds to the C2 barrel (**Fig 9b** *widefield sensory mapping*: pink region) and the best-expressing FOV near this location will be used going forwards (**Fig 9b** *widefield sensory mapping*: white dashed box; **Fig 9b** *Opsin expression* and *Indicator expression*).
7. Allow animal to recover from anaesthesia (hours to days, depending on the experiment)
8. Headfix the awake animal on a spherical treadmill under the microscope and use the 2-photon imaging path to navigate to the best-expressing L2/3 FOV near the C2 barrel (∼150 - 200 µm deep).
9. Map expression (see **Module 3a**, **Fig 5**) by collecting mean 2-photonimages of opsin and indicator expression at their respective wavelengths (GCaMP6s: 920 nm; C1V1-Kv2.1-mRuby: 765 nm). The indicator mean image will be used as the reference image for real-time image registration of 2-photonsensory mapping data and the opsin mean image will be used to identify neurons expressing C1V1 via manual curation.
10. For functional mapping (see **Module 4a**, **Figure 7**) we need to maximise the number of sensory-responsive neurons and ensure that Suite2p has enough data to return high quality ROIs. We will use a collection of whisker stimuli (a moving textured wall, 30 Hz dorso-ventral (DV) and rostro-caudal (RC) piezo vibration of the whole whiskerpad) and we will also acquire 30 minutes of spontaneous activity while the animal sits on the treadmill without whisker stimulation. Wall stimuli consist of a 2 s “move in” period where the wall moves into contact with the whiskers, a 4 s “in place period” and a 2 s “move out” period. Piezo stimuli consist of a 1 s 30 Hz sinusoidal stimulus. All stimuli have a 10 s inter-trial interval.
11. Set up and deliver stimuli while collecting a 2-photon imaging movie for each stimulus type in turn (moving wall, DV piezo, RC piezo; ∼25 minutes), followed by 3 x 10 minute spontaneous activity movies. Note that once each of these movies is set up in PrairieView, run *PrairieLink_RawDataStreamReg* and import the indicator mean image to trigger movie acquisition with real-time registration.
12. While acquiring these movies, import the opsin expression image into Naparm and manually identify all neurons expressing C1V1. Export identified C1V1 centroids (for use in Step 15 below).
13. Once all functional mapping data has been acquired (Step 11), run online Suite2p using the registered movie binary files to return ROIs and traces (∼5 minutes/plane). Use the Suite2p curation GUI to curate ROIs (∼15 minutes).
14. Once complete, use the resulting Suite2p output file (*proc.mat) to save out images of ROIs, ROI centroids and pixel-wise local correlation.
15. Import ROI centroids and C1V1 centroids (Step 12) into Naparm. Select any centroid that appears in either of these two images, but that is not too close to the edge of the FOV where photostimulation efficacy is reduced due to vignetting of the system’s photostimulation path and reduced SLM diffraction efficiency when splitting to extreme angles (if not using galvo hopping).
16. To perform photostimulation mapping, (see **Module 4b**, **Figure 8**) set up a photostimulation mapping protocol where all potential target neurons selected above are photostimulated with parameters similar to those that will be used in the final experiment (in our case 10 x 25 ms spiral stimuli at 20 Hz, 500 ms duration). If the total number of targets exceeds the number that can be targeted simultaneously with sufficient photostimulation laser power per neuron (e.g. 6 mW/neuron and 600 mW total power on sample means an upper limit of 100 neurons simultaneously), split the total number into smaller groups of equal size, each of which is small enough such that all neurons within it are photostimulated with sufficient power. Photostimulate each of these patterns with the stimulus parameters described above as a sequence, changing pattern every 1 s with a sequence repetition interval of 10 s.
17. Run photostimulation mapping protocol while acquiring 2-photon imaging data with real-time registration to generate a Naparm movie (**Module 4b**). Add the registered binary file of this Naparm movie to the list of files used for online Suite2p analysis (i.e. along with the functional mapping data acquired previously) and re-run online Suite2p, following by manual curation.
18. Make stimulus-triggered average traces aligned to the onset of sensory stimuli and photostimulus epochs (**Figure 9b** *Trace STAs*) and analyse post-stimulus periods to identify responsive neurons.
19. Find ROIs that exhibit both strong sensory responses and are reliably photostimulatable (**Figure 9c,d,e**; see **Module 4b, Step 13** for discussion of how to define thresholds). In this case we selected ROIs with ΔF/F > 0.3 in response to at least one sensory stimulus and ΔF/F > 0.3 on ≥ 50% of photostimulation trials. TROUBLESHOOTING
20. Use the centroid locations of these ROIs as target positions for behavioural training and save out associated pixel target images.
21. Import these new target images into TPBS (Two-Photon Behaviour Sequencer; https://github.com/hwpdalgleish/TPBS), along with 2-photontarget images used for training previously (in this case of 200 and 100 random neurons in this FOV). Set desired stimulus power for each pixel target image and associate pixel target images to desired PyBehaviour stimulus types and variations (trial-type). Set number and rate of stimulus repetitions for each trial-type and set trial-type ratios and total number of trials. If desired, set an initial trial buffer of appropriate stimulus type (in our case 10 trials of the easiest trial-type; 2-photonstimulation of 100 random neurons).
22. Once complete, generate the training protocol. This saves out the list of trial-types PyBehaviour will deliver, a folder of the corresponding phasemasks and the XML and GPL that define the stimulus order and position respectively in PrairieView.
23. Import the behaviour and microscope configuration files into relevant software and set up a 2-photon imaging movie of appropriate length in PrairieView. Ensure that PackIO is ready to record all triggers. Put the lickometer in place in front of the mouse. Check that the imaging FOV is still the same as recorded during the mapping phase: pick a bright neuron in the previously acquired mean image and ensure that the pixel location of the centre of this neuron in the current FOV is the same as previous.
24. Begin recording with PackIO, set PrairieView Mark Points ready to receive triggers, set TPBS ready to send out triggers, begin acquiring calcium imaging movie, start PyBehaviour session. TROUBLESHOOTING

**! CAUTION** The total duration of functional and photostimulation mapping followed by a behavioural training session can be long (several hours). Ensure that your protocol accounts for this, i.e. that laser powers used are not damaging the tissue over long time periods, the objective remains sufficiently immersed etc. If necessary, mapping and behaviour can be done on different days if your microscope setup and the preparation are stable enough to be able to navigate back to the exact same FOV in terms of translation, pitch, roll, and yaw. It is also essential to carefully monitor the health and behaviour of experimental animals to ensure that they are not in discomfort or distress during such long periods of head restraint.

## TROUBLESHOOTING

Troubleshooting advice can be found in **Table 1**.

**Table 1.** Troubleshooting table.

| Step | Problem | Possible reason | Solution |
| --- | --- | --- | --- |
| 2-3 | Excessive amount of virus leaks out of injection site during/immediately after injection (not a small amount of “backflow” is acceptable) | Injection speed too fast and/or volume too large. | Reduce injection speed and/or volume. |
| 3a-5 | Indicator is dim | Indicator is under-expressing | Increase the concentration and/or volume of indicator virus injected. |
| 3a-5 | Indicator is very bright, does not appear to change brightness over time and/or fills nuclei | Indicator is over-expressing | Decrease the concentration and/or volume of indicator virus injected. |
| 3a-5 | Opsin reporter is dim | Opsin is under-expressing | Increase the concentration and/or volume of opsin virus injected. |
| 3a-5 | Somatically-restricted opsin expression in distal processes (opsin is no longer somatically restricted) | Opsin is over-expressing | Decrease the concentration and/or volume of opsin virus injected. |
| 3a-5 | Neurons may have small bright punctate regions on processes in neuropil, expression in indicator channel is generally very bright and with expression in both cytosol and nucleus. | Tissue damage from needle penetration or injection | Ensure injection pipette is inserted slowly and carefully. Inject more slowly and/or a smaller volume. Use a thinner, sharper injection pipette. |
| 3a-5 | Both opsin and indicator expression is dim | Injection failed.<br>Microscope misaligned or laser malfunctioning. | Check that aliquots/stocks of indicator/opsin constructs are viable. Avoid excessively diluting constructs and ensure that you inject sufficient volume to see expression. During surgery, ensure that the correct volume has actually been injected (e.g. with a graduated glass pipette). If the correct volume has left injection pipette, observe whether you see virus seeping out of the injection site during/immediately after injection. If virus is leaking out, make sure to<br>effectively penetrate the pia (and dura if still intact), and/or try injecting more slowly and leaving the pipette in place after injection for longer.<br>Check the microscope alignment and that the laser and the collection system are functioning correctly by imaging a known good sample, such as pollen grains or a similar finely structured, highly fluorescent material. |
| 3a-5 | Opsin and indicator do not co-express | Promotor or serotype conflicts | Different combinations of opsin and indicator work better in different brain regions likely due to serotype and promotor affinities, this may require some trial and error. Expressing one of the constructs transgenically, while expressing the other virally, may help. |
| 4a-5 | Poor sensory-evoked responses | Weak indicator expression, suboptimal injection site, suboptimal stimulus parameters, or misalignment of electrical/analysis triggers | The indicator expression may be too low to reliably report the activity evoked by the stimulus, safely increasing the level of expression may help. Ensure stimuli are presented successfully. Ensure all frame pulses and stimulus triggers are being recorded correctly for alignment during stimulus-triggered analysis. Ensure the stimulus is appropriate for the brain region being imaged (it is beneficial to confirm area location through widefield response mapping) |
| 4b-12 | Photostimulation efficacy is poor or failed completely (but expression looks good) | Optical alignment is off, or stimulus parameters are suboptimal | Check alignment of the system (including axial parfocality) and laser power on sample (under objective). Most issues can be found by running the spot burning procedure on a plastic slide. If the optical calibration is correct the issue may be that constructs are expressing well, but levels are too low for successful stimulation/readout. The stimulation parameters (power, duration, number of repeats) may be suboptimal for the preparation. |
| 5-19 | Low number of cells that are both responsive to photostimulation and responsive to the task/sensory stimulus of interest | Expression is low, stimulus is suboptimal | The number of cells that are ultimately addressable is a product of two independent probabilities – that of being responsive to the sensory stimulus, and that of being responsive to the photostimulus. Increasing opsin expression while avoiding signs of ill-health will increase the numbers of photostimulatable cells. Depending on the experimental question the definition of stimulus responsiveness could be relaxed. |
| 5-24 | Animals lick constantly | Animal is too water restricted | Remove the animal from the experimental rig and provide it with water to allow animal to maintain a heavier weight (> 80 %). |
| 5-24 | Animal behaviour is inconsistent and/or they give up easily when transitioning from behaviour boxes to microscope rig | Animals become frustrated or confused | Ensure task design and environment is kept as constant as possible – temperature, sound proofing, white ambient noise. Keep pauses in the training session (e.g. for setting up imaging acquisitions) as short as possible |

## ANTICIPATED RESULTS

Here we have described the key steps involved in designing and executing a successful all-optical experiment. Efficient expression of all-optical constructs should result in hundreds of neurons in a single field-of-view (i.e. 500 µm x 500 µm) which are coexpressing enough opsin and indicator for photostimulation and readout (see **Module 3a, Step 5**) while maintaining cell health. An essential step is to rigorously test the photostimulation response of targeted neurons and factor this into the interpretation of the biological results. We describe a strategy for generating a visually intuitive map of photostimulation responses from all neurons in the desired region in as little as 30 minutes. Using the parameters described here, assuming expression is good and cells are healthy, users should expect to see that >50% of neurons show a reliable, detectible photostimulation response (>0.3 ΔF/F [with GCaMP6s] on >50% of trials; see **Module 4b, Step 13**). This provides a platform for targeted activation of different numbers of neurons, in different spatial and temporal patterns, and examining the resulting effects on simultaneously measured local network activity and on behaviour. Recent work has demonstrated the power of this all-optical strategy in comparing the impact of different ensembles of cells on circuit function and behaviour (Carrillo-Reid *et al*., 2016, 2019; Chettih and Harvey, 2019; Jennings *et al*., 2019; Marshel *et al*., 2019; Russell *et al*., 2019; Dalgleish *et al*., 2020; Robinson *et al*., 2020; Daie *et al*., 2021).

The all-optical approach, while powerful, has several limitations which should be carefully assessed in each experiment. Any experiment requiring the expression of exogenous constructs runs the risk of cytopathology due to excessive expression levels, and this risk is increased in all-optical experiments given the need to express both a calcium buffer (the activity indicator) and membrane channel (the opsin) in the same neurons. Therefore, steps must be taken to monitor and mitigate over-expression while also ensuring sufficient expression for experimental purposes (Tian *et al*., 2009; Chen *et al*., 2013; Packer *et al*., 2015). Users should also be aware of problems affecting animal health and welfare, and the quality of recorded neural activity that can come with using transgenic animals (Steinmetz *et al*., 2017). The specifications of optical systems used for such experiments crucially dictate their experimental strengths and weaknesses. Therefore, we encourage users to rigorously characterise, maintain and report in publications the optical and physiological resolution of the system used for each experiment (e.g. as in (Packer *et al*., 2015; Carrillo-Reid *et al*., 2016; Forli *et al*., 2018; Mardinly *et al*., 2018; Marshel *et al*., 2019)), as well as continuously maintaining any necessary calibrations. This is important for ensuring consistency of results over time and reproducibility between labs, to help with interpretation of results, and to inform experimental design and subsequent analysis pipelines. It should also be noted that all-optical systems, including those described in this protocol, are subject to alignment drift over time and therefore routines should be put in place to ensure that metrics indicating correct functioning are monitored frequently (though this need not always extend to the full rigorous characterisations described above). Photostimulation with two-photon excitation can cause heating and potential photodamage due to various linear and non-linear processes (Podgorski and Ranganathan, 2016; Mardinly *et al*., 2018; Picot *et al*., 2018) and therefore efforts must also be made to mitigate this by using safe levels of laser power throughout experiments and monitoring cell health during the experiment.

Looking forward, the dissemination of the all-optical interrogation approach will crucially depend on the continuous development of more powerful hardware (Mardinly *et al*., 2018; Marshel *et al*., 2019), more intuitive software (this paper and Russell *et al*., 2019), more accurate optical algorithms (Eybposh *et al*., 2020) and more sensitive opsins (Mardinly *et al*., 2018; Marshel *et al*., 2019) and indicators (Dana *et al*., 2019). Beyond these specific avenues for improvement, all-optical interrogation will also benefit from ongoing work to increase our ability to image deep in cortical tissue, through the use of three-photon imaging (Horton *et al*., 2013; Ouzounov *et al*., 2017; Wang *et al*., 2018; Weisenburger *et al*., 2019; Yildirim *et al*., 2019), red-shifted indicators (Zhao *et al*., 2011; Inoue *et al*., 2015; Dana *et al*., 2016), adaptive optics (Wang *et al*., 2015; Sun *et al*., 2016) and GRIN lenses (Levene *et al*., 2004; Jennings *et al*., 2019), as well as approaches which allow us to image more neurons (Tsai *et al*., 2015; Pachitariu *et al*., 2016; Sofroniew *et al*., 2016; Stirman *et al*., 2016; Demas *et al*., 2021) at faster rates (Lu *et al*., 2017; Kazemipour *et al*., 2018; Zhang *et al*., 2019; Wu *et al*., 2020). Finally, the continuous development of genetically encoded voltage indicators (Gong *et al*., 2015; Adam *et al*., 2019; Piatkevich *et al*., 2019; Yu *et al*., 2019) will hopefully pave the way to high resolution all-optical electrophysiology of populations of neurons *in vivo* during behaviour (Lou *et al*., 2016; Adam *et al*., 2019; Fan *et al*., 2020; Adam, 2021).

In summary, the strategy presented in this protocol, building on our efforts in different brain areas (Packer *et al*., 2015; Zhang *et al*., 2018; Russell *et al*., 2019; Dalgleish *et al*., 2020; Robinson *et al*., 2020) as well as those of many other groups (Rickgauer *et al*., 2014; Carrillo-Reid *et al*., 2016, 2019; Shemesh *et al*., 2017; Forli *et al*., 2018; Mardinly *et al*., 2018; Chettih and Harvey, 2019; Jennings *et al*., 2019; Marshel *et al*., 2019; Gill *et al*., 2020), represents a first attempt to provide a standardised protocol for all-optical experiments in any region of the mammalian brain, using a range of hardware. Modifications of this protocol, in particular for surgical procedures tailored to particular preparations, should also allow the protocol to be applied to a wide range of species.

## ACKNOWLEDGEMENTS

We thank Isaac Bianco, Jacques Carolan, and Zihui Zhang for comments on the manuscript; Theshika Jeyaratnam for CA1/CA3 behavioural training and surgeries; Soyon Chun, Agnieszka Jucht and Olivia Houghton for mouse breeding; Selmaan Chettih and Christopher Harvey for developing and sharing the somatically-restricted C1V1 opsin; and Bruker Corporation for technical support. This work was supported by grants from the Wellcome Trust, Gatsby Charitable Foundation, ERC, MRC and the BBSRC.

## REFERENCES

Adam, Y. et al. (2019) ‘Voltage imaging and optogenetics reveal behaviour dependent changes in hippocampal dynamics’, Nature, 569, pp. 413–417.

Adam, Y. (2021) ‘All-optical electrophysiology in behaving animals’, Journal of Neuroscience Methods, 353, p. 109101. doi: 10.1016/j.jneumeth.2021.109101.

Antinucci, P. et al. (2020) ‘A calibrated optogenetic toolbox of stable zebrafish opsin lines’, eLife, 9, pp. 1–31. doi: 10.7554/eLife.54937.

Baker, C. A. et al. (2016) ‘Cellular resolution circuit mapping with temporal-focused excitation of soma-targeted channelrhodopsin’, eLife, 5(AUGUST). doi: 10.7554/eLife.14193.

Bhatia, A., Moza, S. and Bhalla, U. S. (2021) ‘Patterned optogenetic stimulation using a DMD projector’, in Methods in Molecular Biology. Humana Press Inc., pp. 173–188. doi: 10.1007/978-1-0716-0830-2_11.

Boyden, E. S. et al. (2005) ‘Millisecond-timescale, genetically targeted optical control of neural activity’, 8(9), pp. 1263–1268. doi: 10.1038/nn1525.

Carandini, M. and Churchland, A. K. (2013) ‘Probing perceptual decisions in rodents’, Nature Neuroscience. Nat Neurosci, pp. 824–831. doi: 10.1038/nn.3410.

Carrillo-Reid, L. et al. (2016) ‘Imprinting and recalling cortical ensembles’, Science, 353(6300), pp. 691–694. doi: 10.1126/science.aaf7560.

Carrillo-Reid, L. et al. (2019) ‘Controlling Visually Guided Behavior by Holographic Recalling of Cortical Ensembles’, Cell, 178(2). doi: 10.1016/j.cell.2019.05.045.

Chen, T.-W. et al. (2013) ‘Ultrasensitive fluorescent proteins for imaging neuronal activity.’, Nature, 499(7458), pp. 295–300. doi: 10.1038/nature12354.

Chettih, S. N. and Harvey, C. D. (2019) ‘Single-neuron perturbations reveal feature-specific competition in V1’, Nature, 567, pp. 334–340. doi: 10.1038/s41586-019-0997-6.

Daie, K., Svoboda, K. and Druckmann, S. (2021) ‘Targeted photostimulation uncovers circuit motifs supporting short-term memory’, Nature Neuroscience, 24(2), pp. 259–265. doi: 10.1038/s41593-020-00776-3.

Dalgleish, H. W. P. et al. (2020) ‘How many neurons are sufficient for perception of cortical activity?’, eLife, 9, pp. 1–99. doi: 10.7554/eLife.58889.

Dana, H. et al. (2016) ‘Sensitive red protein calcium indicators for imaging neural activity’, eLife, 5(MARCH2016), p. e12727. doi: 10.7554/eLife.12727.

Dana, H. et al. (2019) ‘High-performance calcium sensors for imaging activity in neuronal populations and microcompartments’, Nature Methods, 16(7), pp. 649–657. doi: 10.1038/s41592-019-0435-6.

Demas, J. et al. (2021) ‘High-Speed, Cortex-Wide Volumetric Recording of Neuroactivity at Cellular Resolution using Light Beads Microscopy’, bioRxiv, pp. 1–41.

Denk, W. et al. (1994) ‘Anatomical and functional imaging of neurons using 2-photon laser scanning microscopy’, Journal of Neuroscience Methods, 54(2), pp. 151–162. doi: 10.1016/0165-0270(94)90189-9.

Denk, W., Strickler, J. H. and Webb, W. W. (1990) ‘Two-photon laser scanning fluorescence microscopy’, Science, 248(4951), pp. 73–76. doi: 10.1126/science.2321027.

Dombeck, D. A. et al. (2010) ‘Functional imaging of hippocampal place cells at cellular resolution during virtual navigation’, Nature Neuroscience, 13(11), pp. 1433–1440. doi: 10.1038/nn.2648.

Doron, G. et al. (2014) ‘Spiking Irregularity and Frequency Modulate the Behavioral Report of Single-Neuron Stimulation’, Neuron, 81(3), pp. 653–663. doi: 10.1016/j.neuron.2013.11.032.

Eybposh, M. H. et al. (2020) ‘DeepCGH: 3D computer-generated holography using deep learning’, Optics Express, 28(18), p. 26636. doi: 10.1364/oe.399624.

Fan, L. Z. et al. (2020) ‘All-Optical Electrophysiology Reveals the Role of Lateral Inhibition in Sensory Processing in Cortical Layer 1’, Cell, 180, pp. 1–15. doi: 10.1016/j.cell.2020.01.001.

Forli, A. et al. (2018) ‘Two-Photon Bidirectional Control and Imaging of Neuronal Excitability with High Spatial Resolution In Vivo’, Cell Reports, 22(11), pp. 3087–3098. doi: 10.1016/j.celrep.2018.02.063.

Forli, A. et al. (2021) ‘Optogenetic strategies for high-efficiency all-optical interrogation using blue light-sensitive opsins’, eLife, 10. doi: 10.7554/eLife.63359.

Gerchberg, R. W. and Saxton, W. O. (1972) ‘PRACTICAL ALGORITHM FOR THE DETERMINATION OF PHASE FROM IMAGE AND DIFFRACTION PLANE PICTURES.’, Optik (Stuttgart), 35(2), pp. 237–250.

Gill, J. V. et al. (2020) ‘Precise Holographic Manipulation of Olfactory Circuits Reveals Coding Features Determining Perceptual Detection’, Neuron, 108(2), pp. 382–393.e5. doi: 10.1016/j.neuron.2020.07.034.

Giovannucci, A. et al. (2019) ‘CaImAn an open source tool for scalable calcium imaging data analysis’, eLife, 8. doi: 10.7554/eLife.38173.

Gollisch, T. and Meister, M. (2008) ‘Rapid Neural Coding in the Retina with Relative Spike Latencies’, Science, 319(February), pp. 1108–1112.

Gong, Y. et al. (2015) ‘High-speed recording of neural spikes in awake mice and flies with a fluorescent voltage sensor’, Science, 350(6266), pp. 1361–1366. doi: 10.1126/science.aab0810.

Grienberger, C. and Konnerth, A. (2012) ‘Imaging calcium in neurons.’, Neuron, 73(5), pp. 862–85. doi: 10.1016/j.neuron.2012.02.011.

Grosenick, L., Marshel, J. H. and Deisseroth, K. (2015) ‘Closed-loop and activity-guided optogenetic control’, Neuron. Cell Press, pp. 106–139. doi: 10.1016/j.neuron.2015.03.034.

Guo, Z. V. et al. (2014) ‘Procedures for Behavioral Experiments in Head-Fixed Mice’, PLoS ONE. Edited by S. A. Simon, 9(2), p. e88678. doi: 10.1371/journal.pone.0088678.

Harris, K. D. and Mrsic-Flogel, T. D. (2013) ‘Cortical connectivity and sensory coding’, Nature. Nature Publishing Group, pp. 51–58. doi: 10.1038/nature12654.

Histed, M. H. and Maunsell, J. H. R. (2014) ‘Cortical neural populations can guide behavior by integrating inputs linearly, independent of synchrony.’, Proceedings of the National Academy of Sciences, 111(1), pp. E178–87. doi: 10.1073/pnas.1318750111.

Holtmaat, A. et al. (2009) ‘Long-term, high-resolution imaging in the mouse neocortex through a chronic cranial window’, Nature Protocols, 4(8), pp. 1128–1144. doi: 10.1038/nprot.2009.89.

Hopt, A. and Neher, E. (2001) ‘Highly nonlinear photodamage in two-photon fluorescence microscopy’, Biophysical Journal, 80(4), pp. 2029–2036. doi: 10.1016/S0006-3495(01)76173-5.

Horton, N. G. et al. (2013) ‘In vivo three-photon microscopy of subcortical structures within an intact mouse brain’, Nature Photonics, 7(3), pp. 205–209. doi: 10.1038/nphoton.2012.336.

Huber, D. et al. (2008) ‘Sparse optical microstimulation in barrel cortex drives learned behaviour in freely moving mice.’, Nature, 451(7174), pp. 61–4. doi: 10.1038/nature06445.

Inoue, M. et al. (2015) ‘Rational design of a high-affinity, fast, red calcium indicator R-CaMP2’, Nature Methods, 12(1), pp. 64–70. doi: 10.1038/nmeth.3185.

Inoue, M. et al. (2019) ‘Rational Engineering of XCaMPs, a Multicolor GECI Suite for In Vivo Imaging of Complex Brain Circuit Dynamics’, Cell, 177(5), pp. 1346–1360.e24. doi: 10.1016/j.cell.2019.04.007.

Jacobs, A. L. et al. (2009) ‘Ruling out and ruling in neural codes’, Proceedings of the National Academy of Sciences of the United States of America, 106(14), pp. 5936–5941.

Jennings, J. H. et al. (2019) ‘Interacting neural ensembles in orbitofrontal cortex for social and feeding behaviour’, Nature 2019, p. 1. doi: 10.1038/s41586-018-0866-8.

Kazemipour, A. et al. (2018) ‘Kilohertz frame-rate two-photon tomography’, bioRxiv.

Klapoetke, N. C. et al. (2014) ‘Independent optical excitation of distinct neural populations’, Nature Methods, 11(3), pp. 338–346. doi: 10.1038/nmeth.2836.

Koester, H. J. et al. (1999) ‘Ca2+ fluorescence imaging with pico- and femtosecond two-photon excitation: Signal and photodamage’, Biophysical Journal, 77(4), pp. 2226–2236. doi: 10.1016/S0006-3495(99)77063-3.

Levene, M. J. et al. (2004) ‘In Vivo Multiphoton Microscopy of Deep Brain Tissue’, Journal of Neurophysiology, 91(4), pp. 1908–1912. doi: 10.1152/jn.01007.2003.

Lim, S. T. et al. (2000) ‘A novel targeting signal for proximal clustering of the Kv2.1 K+ channel in hippocampal neurons’, Neuron, 25(2), pp. 385–397. doi: 10.1016/S0896-6273(00)80902-2.

London, M. et al. (2010) ‘Sensitivity to perturbations in vivo implies high noise and suggests rate coding in cortex.’, Nature, 466(7302), pp. 123–127. doi: 10.1038/nature09086.

Lou, S. et al. (2016) ‘Genetically targeted all-optical electrophysiology with a transgenic cre-dependent optopatch mouse’, Journal of Neuroscience, 36(43), pp. 11059–11073. doi: 10.1523/JNEUROSCI.1582-16.2016.

Lu, R. et al. (2017) ‘Video-rate volumetric functional imaging of the brain at synaptic resolution’, Nature Neuroscience, 20(4), pp. 620–628. doi: 10.1038/nn.4516.

Mardinly, A. R. et al. (2018) ‘Precise multimodal optical control of neural ensemble activity’, Nature Neuroscience. doi: 10.1038/s41593-018-0139-8.

Marshel, J. H. et al. (2019) ‘Cortical layer–specific critical dynamics triggering perception’, Science, 365, pp. 1–12. doi: 10.1126/science.aaw5202.

Mattis, J. et al. (2012) ‘Principles for applying optogenetic tools derived from direct comparative analysis of microbial opsins.’, Nature Methods, 9(2), pp. 159–172. doi: 10.1038/nmeth.1808.

Nikolenko, V. et al. (2008) ‘SLM microscopy: scanless two-photon imaging and photostimulation with spatial light modulators’, Frontiers in Neural Circuits, 2(December), pp. 1–14. doi: 10.3389/neuro.04.005.2008.

Ouzounov, D. G. et al. (2017) ‘In vivo three-photon imaging of activity of GCaMP6-labeled neurons deep in intact mouse brain’, Nature Methods, 14(4), pp. 388–390. doi: 10.1038/nmeth.4183.

Pachitariu, M. et al. (2016) ‘Suite2p: beyond 10,000 neurons with standard two-photon microscopy’, bioRxiv, p. 061507. doi: 10.1101/061507.

Packer, A. M. et al. (2012) ‘Two-photon optogenetics of dendritic spines and neural circuits’, Nature Methods, 9(12), pp. 1202–1208. doi: 10.1038/NMETH.2249.

Packer, A. M. et al. (2015) ‘Simultaneous all-optical manipulation and recording of neural circuit activity with cellular resolution in vivo’, Nature Methods, 12(2). doi: 10.1038/nmeth.3217.

Panzeri, S. et al. (2001) ‘The role of spike timing in the coding of stimulus location in rat somatosensory cortex.’, Neuron, 29(3), pp. 769–77.

Panzeri, S. et al. (2017) ‘Cracking the neural code for sensory perception by combining statistics, intervention and behavior’, Neuron, 93(3), pp. 491–507. doi: 10.1111/gcb.13051.

Papagiakoumou, E. et al. (2008) ‘Patterned two-photon illumination by spatiotemporal shaping of ultrashort pulses’, Optics Express, 16(26), p. 22039. doi: 10.1364/oe.16.022039.

Papagiakoumou, E. et al. (2010) ‘Scanless two-photon excitation of channelrhodopsin-2’, Nature Methods, 7(10), pp. 848–854. doi: 10.1038/nmeth.1505.

Papagiakoumou, E., Ronzitti, E. and Emiliani, V. (2020) ‘Scanless two-photon excitation with temporal focusing’, Nature Methods. Nature Research, pp. 571–581. doi: 10.1038/s41592-020-0795-y.

Pégard, N. C. et al. (2017) ‘Three-dimensional scanless holographic optogenetics with temporal focusing (3D-SHOT)’, Nature Communications, 8(1), pp. 1–14. doi: 10.1038/s41467-017-01031-3.

Peron, S. et al. (2020) ‘Recurrent interactions in local cortical circuits’, Nature, 579(7798), pp. 256–259. doi: 10.1038/s41586-020-2062-x.

Peron, S. and Svoboda, K. (2011) ‘From cudgel to scalpel: toward precise neural control with optogenetics’, Nature Methods, 8(1), pp. 30–34. doi: 10.1038/NMETH.F.325.

Piatkevich, K. D. et al. (2019) ‘Population imaging of neural activity in awake behaving mice’, Nature, 574, pp. 413–417. doi: 10.1038/s41586-019-1641-1.

Picot, A. et al. (2018) ‘Temperature Rise under Two-Photon Optogenetic Brain Stimulation’, Cell Reports, 24(5), pp. 1243–1253.e5. doi: 10.1016/j.celrep.2018.06.119.

Pinto, L. and Dan, Y. (2015) ‘Cell-Type-Specific Activity in Prefrontal Cortex during Goal-Directed Behavior’, Neuron, 87(2), pp. 437–450. doi: 10.1016/j.neuron.2015.06.021.

Podgorski, K. and Ranganathan, G. (2016) ‘Brain heating induced by near-infrared lasers during multiphoton microscopy’, Journal of Neurophysiology, 116(3), pp. 1012–1023. doi: 10.1152/jn.00275.2016.

Prakash, R. et al. (2012) ‘Two-photon optogenetic toolbox for fast inhibition, excitation and bistable modulation’, Nature Methods, 9(12), pp. 5–7. doi: 10.1038/NMETH.2215.

Rickgauer, J. P., Deisseroth, K. and Tank, D. W. (2014) ‘Simultaneous cellular-resolution optical perturbation and imaging of place cell firing fields’, Nature Neuroscience, 17(12). doi: 10.1038/nn.3866.

Rickgauer, J. P. and Tank, D. W. (2009) ‘Two-photon excitation of channelrhodopsin-2 at saturation.’, Proceedings of the National Academy of Sciences, 106(35), pp. 15025–15030. doi: 10.1073/pnas.0907084106.

Rickgauer, J. and Tank, D. (2009) ‘Two-Photon Excitation of Channelrhodopsin-2 at Saturation’, Proceedings of the National Academy of Sciences of the United States of America, 106(35), pp. 15025–15030. doi: 10.1038/nmeth.1505.

Robinson, N. T. M. et al. (2020) ‘Targeted Activation of Hippocampal Place Cells Drives Memory-Guided Spatial Behavior.’, Cell, 183(7), pp. 2041–2042. doi: 10.1016/j.cell.2020.12.010.

Russell, L. E. et al. (2019) ‘The influence of visual cortex on perception is modulated by behavioural state’, bioRxiv. bioRxiv, p. 706010. doi: 10.1101/706010.

Shemesh, O. A. et al. (2017) ‘Temporally precise single-cell-resolution optogenetics’, Nature Neuroscience, 20(12), pp. 1796–1806. doi: 10.1038/s41593-017-0018-8.

Shusterman, R. et al. (2011) ‘Precise olfactory responses tile the sniff cycle’, Nature Neuroscience, 14(8), pp. 1039–1044. doi: 10.1038/nn.2877.

Sofroniew, N. J. et al. (2016) ‘A large field of view two-photon mesoscope with subcellular resolution for in vivo imaging’, eLife, 5, p. e14472. doi: 10.7554/eLife.14472.

Sridharan, S. et al. (2021) ‘High performance microbial opsins for spatially and temporally precise perturbations of large neuronal networks’, bioRxiv.

Steinmetz, N. A. et al. (2017) ‘Aberrant Cortical Activity In Multiple GCaMP6-Expressing Transgenic Mouse Lines’, eNeuro, 4(October), pp. 1–15. doi: 10.1101/138511.

Stirman, J. N. et al. (2016) ‘Wide field-of-view, multi-region, two-photon imaging of neuronal activity in the mammalian brain’, Nature Biotechnology, 34(8), pp. 865–870. doi: 10.1038/nbt.3594.

Sun, W. et al. (2016) ‘Thalamus provides layer 4 of primary visual cortex with orientation- and direction-tuned inputs’, Nature Neuroscience, 19(2), pp. 308–315. doi: 10.1038/nn.4196.

Szabo, V. et al. (2014) ‘Spatially selective holographic photoactivation and functional fluorescence imaging in freely behaving mice with a fiberscope’, Neuron, 84(6), pp. 1157– 1169. doi: 10.1016/j.neuron.2014.11.005.

Tian, L. et al. (2009) ‘Imaging neural activity in worms, flies and mice with improved GCaMP calcium indicators’, Nature Methods, 6(12), pp. 875–881. doi: 10.1038/nmeth.1398.

Tsai, P. S. et al. (2015) ‘Ultra-large field-of-view two-photon microscopy’, Optics Express, 23(11), p. 13833. doi: 10.1364/oe.23.013833.

Wang, K. et al. (2015) ‘Direct wavefront sensing for high-resolution in vivo imaging in scattering tissue’, Nature Communications, 6, pp. 1–6. doi: 10.1038/ncomms8276.

Wang, T. et al. (2018) ‘Three-photon imaging of mouse brain structure and function through the intact skull’, Nature Methods, 15(10), pp. 789–792. doi: 10.1038/s41592-018-0115-y.

Weisenburger, S. et al. (2019) ‘Volumetric Ca2+ Imaging in the Mouse Brain Using Hybrid Multiplexed Sculpted Light Microscopy’, Cell, 177(4). doi: 10.1016/j.cell.2019.03.011.

Wiegert, J. S. et al. (2017) ‘Silencing Neurons: Tools, Applications, and Experimental Constraints’, Neuron, 95(3), pp. 504–529. doi: 10.1016/j.neuron.2017.06.050.

Wu, J. et al. (2020) ‘Kilohertz two-photon fluorescence microscopy imaging of neural activity in vivo’, Nature Methods, 17(March), pp. 287–290. doi: 10.1038/s41592-020-0762-7.

Yang, W. et al. (2018) ‘Simultaneous two-photon imaging and two-photon optogenetics of cortical circuits in three dimensions’, eLife, 7. doi: 10.7554/eLife.32671.

Yildirim, M. et al. (2019) ‘Functional imaging of visual cortical layers and subplate in awake mice with optimized three-photon microscopy’, Nature Communications, 10(177), pp. 1–12. doi: 10.1038/s41467-018-08179-6.

Yizhar, O., Fenno, Lief E., et al. (2011) ‘Neocortical excitation/inhibition balance in information processing and social dysfunction’, Nature, 477(7363), pp. 171–178. doi: 10.1038/nature10360.

Yizhar, O., Fenno, Lief E, et al. (2011) ‘Optogenetics in Neural Systems’, Neuron, 71(1), pp. 9–34. doi: 10.1016/j.neuron.2011.06.004.

Yu, J. et al. (2019) ‘Bright and photostable chemigenetic indicators for extended in vivo voltage imaging’, Science, 704(August), pp. 699–704.

Zhang, F. et al. (2011) ‘The microbial opsin family of optogenetic tools’, Cell. Cell Press, pp. 1446–1457. doi: 10.1016/j.cell.2011.12.004.

Zhang, T. et al. (2019) ‘Kilohertz two-photon brain imaging in awake mice’, Nature Methods, 16(November). doi: 10.1038/s41592-019-0597-2.

Zhang, Z. et al. (2018) ‘Closed-loop all-optical interrogation of neural circuits in vivo’, Nature Methods, 15(December), pp. 1037–1040. doi: 10.1038/s41592-018-0183-z.

Zhao, Y. et al. (2011) ‘An Expanded Palette of Genetically Encoded Ca2+ Indicators’, Cell, 557(September), pp. 1888–1891.

